# An experimental and theoretical approach to understand the interaction between particles and mucosal tissues

**DOI:** 10.1101/2022.09.15.508137

**Authors:** Roni Sverdlov Arzi, Maya Davidovich-Pinhas, Noy Cohen, Alejandro Sosnik

## Abstract

Nanonization of poorly water-soluble drugs has shown great potential in improving their oral bioavailability by enhancing the dissolution rate and saturation solubility. Moreover, due to particle size reduction and larger surface area, the number of contact points with the gastrointestinal mucus favors adhesion. Similar phenomena could be anticipated when nano-pollutants come into direct contact with mucosal tissues. However, the fundamental features that govern the interaction of particles with mucus have not been investigated in a systematic and rational way before. In this work, we synthesize mucin hydrogels of different pore sizes with rheological properties that closely mimic the properties of freshly extracted porcine mucin. By using fluorescent pure curcumin particles, we characterize the effect of particle size (hydrodynamic diameter of 200 nm, and 1.2 and 1.3 μm), concentration (18, 35, and 71 μg mL^−1^), and hydrogel crosslinking density (which is directly related to the stiffness and governs the average pore size) on the diffusion-driven particle penetration *in vitro*. Next, we derive a phenomenological model that describes the physics behind the diffusion-derived penetration of particles into the mucin network and considers the contributions of the particle size, the particle concentration, and the crosslinking density of the mucin hydrogel. Finally, we challenge our experimental-theoretical approach by preliminarily assessing the oral pharmacokinetics of an anti-cancer model drug, namely dasatinib, in pristine and nanonized forms and two clinically relevant doses in rats. For of a dose of 10 mg kg^−1^, drug nanonization leads to a significant ~8- and ~21-fold increase of the drug oral bioavailability and half-life, respectively, with respect to the unprocessed micron-sized drug. When the drug dose of pure drug nanoparticles (which is directly related to the local concentration of the drug in the gastrointestinal tract) was increased to 15 mg mL^−1^, the oral bioavailability increased though not significantly, suggesting the saturation of the penetration sites in the mucus, as demonstrated by the *in vitro* model. Our overall results reveal the potential of this experimental-theoretical approach, shed light on the interaction of particulate matter and mucosal tissues, and pave the way for the development of tools that enable a more rational design of nano-drug delivery systems for mucosal administration and the assessment of risk factors related to the exposure of mucosal tissues to nano-pollutants.

## 1. Introduction

Mucosal tissues (e.g., gastrointestinal tract, airways, eyes, urinary and genital systems) constitute a major interface between the human body and the external milieu and they perform as barriers that control the permeability of exogenous soluble, supramolecular and particulate matter [1], while permitting the exchange of water, gases and other essential nutrients [2]. Depending on the body site and the physiological function, mucosae exhibit different cellular structure [3]. At the same time, they share a common feature: they are all covered by a dynamic layer of mucus which is a water-rich (>90% water) non-Newtonian porous mucin gel with a viscosity in the range of 10—10^3^ Pa·s and variable thickness that protects the underlying tissue from external physicochemical, mechanical and biological insults [4–7]. Mucins are high molecular weight glycoproteins formed by a central protein core and highly glycosylated “bottle-brush”-like and “naked” regions [8,9] and, upon hydration, mucin fibers form a dense and hierarchical network of pores with diameters between few tens and several hundreds of nanometers [8,10]. Another common feature of mucosal tissues is that they host components of the immune system such as naïve, activated and memory B cells, T cells, dendritic cells and macrophages in the so-called mucosa-associated lymphoid tissue (MALT) which is a compartmentalized system that functions independently of the systemic immunity [3,11,12].

The unique structure and properties of mucus have been capitalized on to design mucoadhesive and muco-penetrating drug delivery systems that improve the local and trans-mucosal delivery of a broad spectrum of active agents, from small-molecule drugs to macromolecules [13–15]. The specific surface area-to-volume and number of contact points of nanomaterials with the surrounding tissues is dramatically greater than that of large particles and macroscopic formulations (e.g., tablets) and it also leads to changes of the intrinsic thermodynamic and kinetic properties of the raw material [16,17]. Over the last decades, particle size reduction techniques such as micronization and nanonization have gained clinical impact [16,18,19]. Pure drug nanoparticles (PDNPs) are the simplest and most patented technology to increase the dissolution rate of hydrophobic drugs in biological fluids [20]. PDNPs can be dispersed in aqueous media (i.e., nanosuspensions) [19,21] or used to produce solid formulations and, as opposed to other drug nanocarriers (i.e., liposomes, polymeric nanoparticles) for which encapsulation efficiency and drug loading have to be defined in the final product, PDNPs offer a theoretical drug content of up to 100% which improves their chances of bench-to-bedside translation [22,23]. In oral drug delivery, PDNPs have been mainly envisioned to increase the dissolution rate and the aqueous saturation solubility of poorly water-soluble drugs in the gastrointestinal tract (GIT) and their oral bioavailability which is often evidenced by an increase of the maximum concentration (C_max_) and the area-under-the-curve (AUC) in the pharmacokinetic curve (plasma concentration versus time) [16,22,23]. Intriguingly, we showed in a previous work that nanonization of the hydrophobic protease inhibitor antiretroviral indinavir free base not only significantly increased the C_max_ and the AUC by 4- and 22-fold, respectively, but also resulted in a dramatic prolongation of the half-life (t_1/2_) from 0.8 to 22 h with respect to the raw drug, a phenomenon that could be explained only by the entrapment of pure indinavir nanoparticles in the intestinal mucus and their slow dissolution and absorption over time [24]. Since mucus is a hierarchical porous hydrated negatively-charged matrix, it is anticipated that it can discriminate among particles by size, density, surface charge and chemistry, and shape [25]. However, the fundamental features that govern the interaction of particles with mucus have not been investigated in a systematic and rational way before. Furthermore, attempts to anticipate the fate of those interactions based on known particle and mucus features, and mucoadhesion theories [26] can be misleading due to the great variability of the mucus in different body sites and even along portions of the same system (i.e., the gut), and under various medical conditions [15,27]. A fundamental study of this kind would be crucial to rationalize the design of different nano-drug delivery systems for mucosal administration and predict their performance *in vivo*, which may reduce the drug attrition rates [28]. The investigation of these interactions is also crucial to understand the toxicological implications of the exposure of the human body to nano-pollutants released into the air and water effluents by accident or as byproducts of industrial processes and that do not undergo proper decontamination [29]. Nano-pollutants come into direct acute or chronic contact with mucosal tissues (especially the airways and the GIT), in which they can penetrate the mucus layer and trigger a local immune response mediated by the MALT and thus, lead to local toxicity (i.e., asbestos) or translocate the epithelial layer, reach the bloodstream and cause systemic toxicity [30–33]. The experimental-theoretical tools available to model the interactions between nanoparticulate matter and mucus are extremely limited and novel approaches are called for [34].

Aiming to gain further insight into the penetration mechanisms that govern particle penetration and entrapment, in a previous work we derived a preliminary methodical statistical-mechanics based model that describes the local interactions between particles and biological gels (i.e., mucus) upon spontaneous (diffusion-driven) and forced penetration [35]. The model resulted in relations that enable the quantitative measurement of the changes in the network due to particle penetration and included the derivation of the relationship between (i) the external force exerted on the network and (ii) the stretch of the polymeric chains in the hydrogel, and the particle sink. Nevertheless, our previous work assumed that there are no interactions between adjacent particles (i.e., low particle concentration) and did not investigate the influence of fundamental aspects such as particle size and concentration, and changes in the porosity of the gel on these interactions.

In this work, we investigate and develop an experimental-theoretical approach to predict and understand the interaction of particles with mucosal tissues. To study the effect of key physicochemical parameters on the penetration process, we synthesize mucin hydrogels that mimic the rheological and structural properties of mucus in the GIT and characterize the effect of particle size and concentration and hydrogel crosslinking density (which directly affects pore size and viscoelasticity) on particle penetration under diffusion-driven penetration *in vitro*. Finally, we challenge these findings in a preliminary study of the oral pharmacokinetics of a model anti-cancer poorly water-soluble drug in rats.

## 2. Results and Discussion

### 2.1. Physical aspects of particle penetration into hydrogels

The penetration of particulate matter into a hydrogel network such as mucus is enabled by the intermolecular spacing between polymer chains (also referred to as mesh-size), which is proportional to the average distance between crosslinks [36,37]. These spacings set a threshold beyond which the transport of particles is reduced or completely inhibited owing to steric hindrance [15,35,38]. Specifically, particles with dimensions smaller than the interchain spacing that do not chemically interact with the hydrogel are expected to penetrate the network (**Figure 1A**). Such mechanism is referred to as unrestricted diffusion. Conversely, particles that are larger than the local spacings must be forced into the network, leading to the local stretching of polymer chains. The increase in the intermolecular spacing enables particle penetration, as shown in **Figure 1B**. This mechanism is henceforth referred to as restricted diffusion. Examples of forces that are externally exerted on the mucus layer of mucosal tissues under physiological conditions include, for example, peristalsis in the GIT, the blinking of the eyes, and coughing [39].

**Figure 1.**
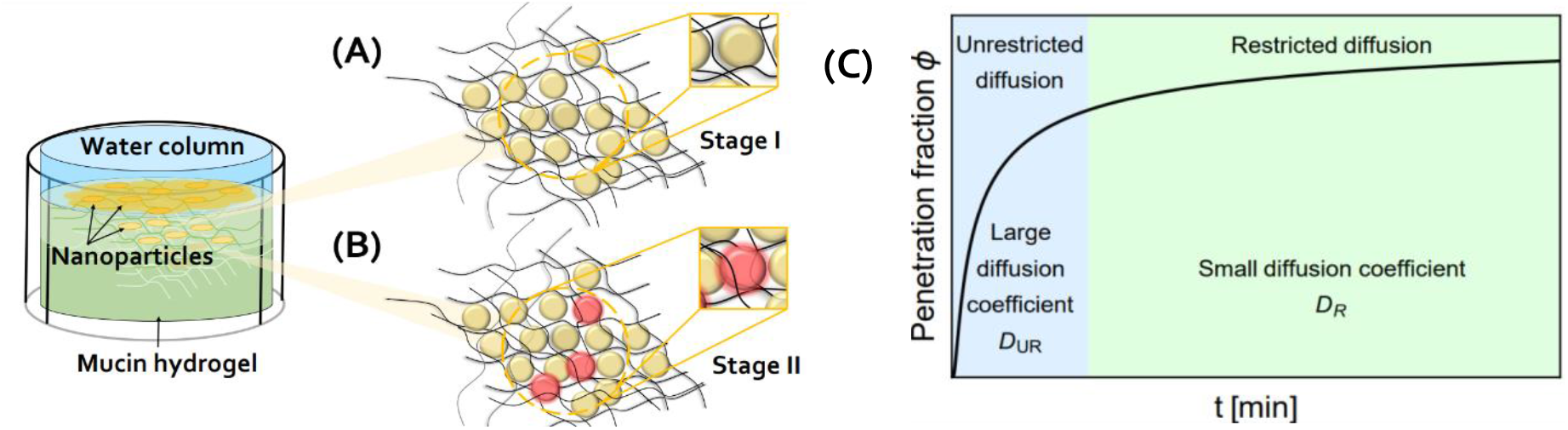
Particle penetration into a hydrogel. Scheme of (A) unrestricted and (B) restricted diffusion of particles into a hydrogel network. (C) Illustration of the time-evolution of the overall fraction of particles in the hydrogel.

The diffusion-driven penetration of particles into the mucin hydrogel can be viewed as a two-stage process [40–42], as described in **Figure 1C**. In the first stage, particles unrestrictedly diffuse into areas along the mucin hydrogel surface in which the intermolecular spacing is larger than their diameter. Accordingly, the diffusion coefficient in this stage can be estimated *via* the Stokes-Einstein equation,

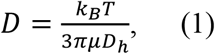

which captures the movement of particles through a continuous liquid [40–42]. Here, *k_B_* is the Boltzmann constant, *T* denotes the temperature, *μ* is the medium viscosity, and *D_h_* is the average diameter of the particles. If we consider the penetration of particles with a diameter range of several hundreds of nanometers up to ~1 μm in water at room temperature (RT, *μ* ~ 10^−3^ *Pa* · *s*), the calculated diffusion coefficients would be in the range of *D* ~ 0.3 — 2.2 μm s^−1^.

In the second stage, the diffusion-driven penetration of additional particles becomes restricted and requires the local stretching of polymeric chains to increase the intermolecular spacings that are originally smaller than the particle diameter (**Figure 1B**). The diffusion coefficient in the second stage depends on the characteristics of the particles and the network (examples include particle diameter, concentration, and crosslinking density of the network [43,44]). It is underscored that the diffusion coefficient is significantly reduced upon the transition from stage I to stage II (**Figure 1C**).

To model the overall diffusion (in both the unrestricted and the restricted stages), we propose the relation

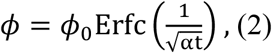

where *ϕ*_0_ is the maximum fraction of particles that penetrate the mucin hydrogel (i.e., the percentage of particles in the hydrogel at steady state) and α is a constant that accounts for the effective diffusion coefficient of the particles in the two stages and the geometry of the gel. Since the diffusion process depends on the particles and the network, we further propose to multiplicatively decompose *ϕ*_0_ into the three contributions that account for the particle size ϕ_D_h__(D_h_), the particle concentration ϕ_C_(C), and crosslinking density of the mucin hydrogel Φ_G’_(G’), such that

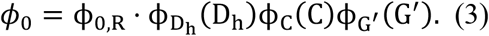

Here, ϕ_0,R_ is the value of ϕ_0_ for a reference system with characteristic particle size, particle concentration, and hydrogel crosslinking density, as detailed below. The contribution of each factor assumes the form

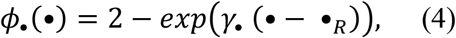

where • = D_h_, C, G’, *γ*_•_ expresses the sensitivity of the system to the change in the • parameter, and •_R_ is the reference value of •. Note that for the baseline values of the reference system *ϕ*_•_(•) = 1 and thus, *ϕ*_0_ = ϕ_0,R_.

Next, we propose an experimental design to assess the penetration of particulate matter into mucin hydrogels that mimic the intestinal mucus.

### 2.2. Particle penetration into mucin hydrogels *in vitro*

#### 2.2.1. Synthesis and characterization of mucus-mimicking mucin hydrogels

To produce mucin hydrogels that mimic the viscoelastic and structural features of gastrointestinal mucus, we exhaustively methacrylated porcine gastric mucin by reacting pendant primary amine moieties in the glycoprotein with methacrylic anhydride (MA) at a 1:2 mucin:MA weight ratio in water under light basic conditions (pH 8.0) [45] and 4°C for 12 h [46], as depicted in **Figure 2**. The degree of methacrylation of methacrylated mucin (mucin-MA) determines the maximal crosslinking density achievable upon free radical polymerization, which affects the mesh size and the stiffness of the mucin hydrogel. When fully crosslinked, a hydrogel with a high degree of methacrylation will display higher stiffness and smaller mesh size than a counterpart with low degree of methacrylation and vice versa [47]. The methacrylation efficiency was determined by the 2,4,6-trinitrobenzenesulfonic acid (TNBSA) assay. TNBSA is a reagent that reacts with the free amine groups that do not undergo methacrylation, giving an indirect indication on the number of groups that reacted and thus, upon crosslinking, can potentially form crosslinking sites [48]. The methacrylation efficiency was 99.7 ± 1%, indicating that almost all the available sites for methacrylation in mucin and later crosslinking were chemically modified.

**Figure 2.**
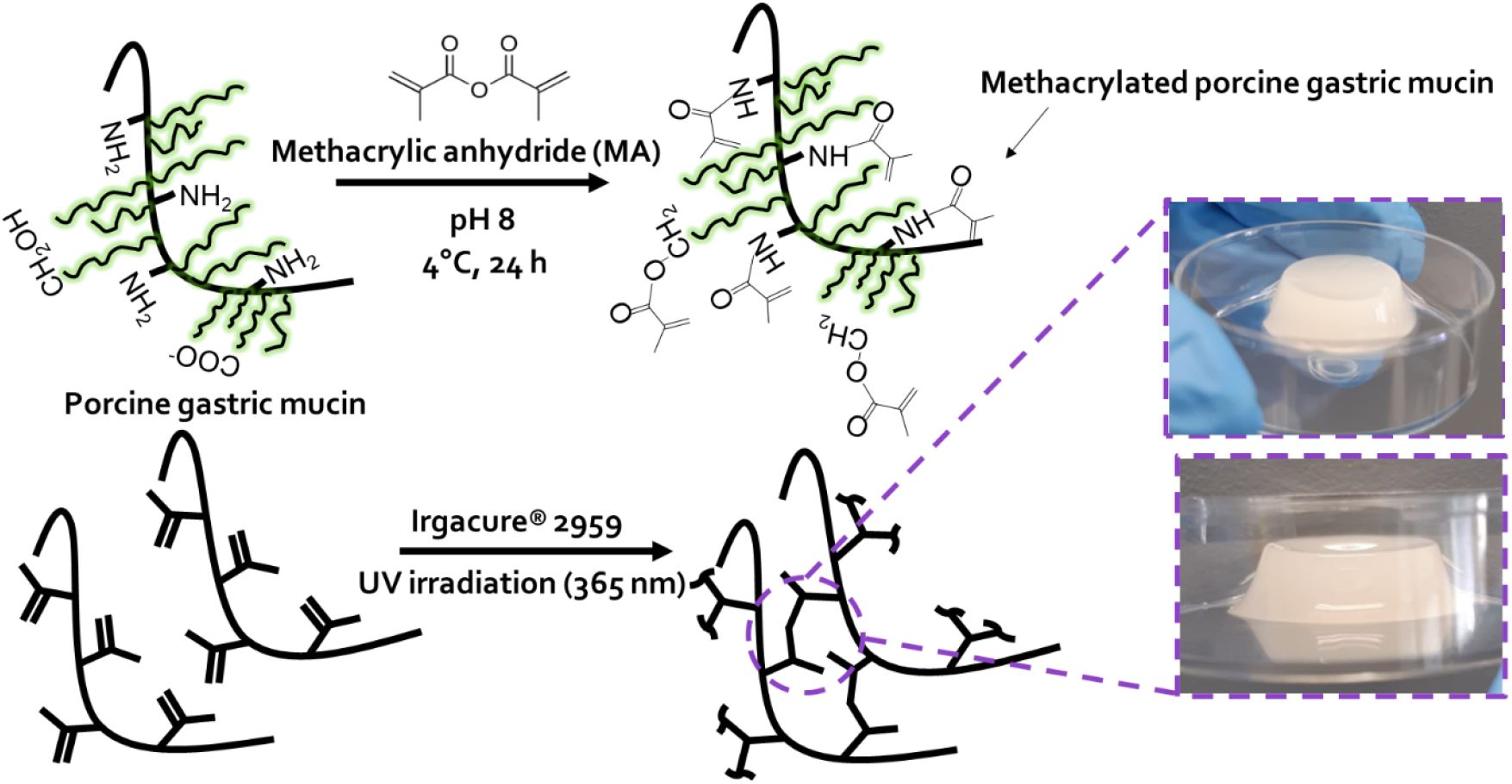
Synthesis of mucus-mimicking mucin hydrogels *via* photo-initiated crosslinking. Porcine gastric mucin was methacrylated by reaction with methacrylic anhydride in water and then, crosslinked using Irgacure^®^ 2959 as photo-initiator and UV light (λ = 365 nm).

To prepare mucus-mimicking hydrogels, mucin-MA was dissolved in water (40 mg mL^−1^), poured into 24-well plates (1 mL) and photo-crosslinked under UV light (365 nm, 8 min) using 2-hydroxy-4-(2-hydroxyethoxy)-2-methylpropiophenone (Irgacure^®^ 2959) (**Figure 2**), a highly efficient radical Type I photo-initiator for UV curing systems of unsaturated monomers and/or prepolymers [49–52]. Unlike other UV photo-initiators, it has shown very low cytotoxicity, making it suitable for use in biomedical applications such as tissue engineering, even in the presence of cells during the crosslinking stage [53,54]. Since Irgacure^®^ 2959 displays limited solubility in water (≤0.5% w/v at RT), we dissolved it in ethanol prior to the addition of a small volume of a concentrated ethanolic solution to the mucin-MA solution [53–56].

To synthesize hydrogels of variable crosslinking density, we could modify the degree of methacrylation or, conversely, adjust the photo-initiator concentration. We found out that an increase of the final photo-initiator concentration in the precursor mucin-MA solution resulted in softer hydrogels because the crosslinking efficiency of Irgacure^®^ 2959 under UV light depends on its final concentration in the precursor solution and also on the relative concentration of ethanol (or any other water-miscible organic solvent) as solvent traces reduce the efficiency of the free radical polymerization reaction [57]. In this work, we used a photo-initiator solution with a constant concentration of 10 mg mL^−1^ in ethanol and thus, the higher its final concentration in the aqueous mucin-MA solution, the greater the ethanol volume added to it, enabling the fine-tuning of the degree of crosslinking. Based on this, lightly-crosslinked (L-CL), moderately-crosslinked (M-CL), and highly-crosslinked (H-CL) mucin hydrogels were obtained. The diameter and the thickness of the hydrogels were 15.6 and 8.5 mm, respectively. The rheological properties of L-CL, M-CL, and H-CL mucin hydrogels were characterized in a plate rheometer on hydrogels swollen in water and left to equilibrate for at least 24 h before the test. First, a strain sweep at a constant frequency of 10 rad s^−1^ was performed to extract the linear viscoelastic (LVE) range of the hydrogels, in which the test can be carried out without disturbing the hydrogel structure (**Figure 3A**), in compliance with the principle of small deformation rheology [58].

**Figure 3.**
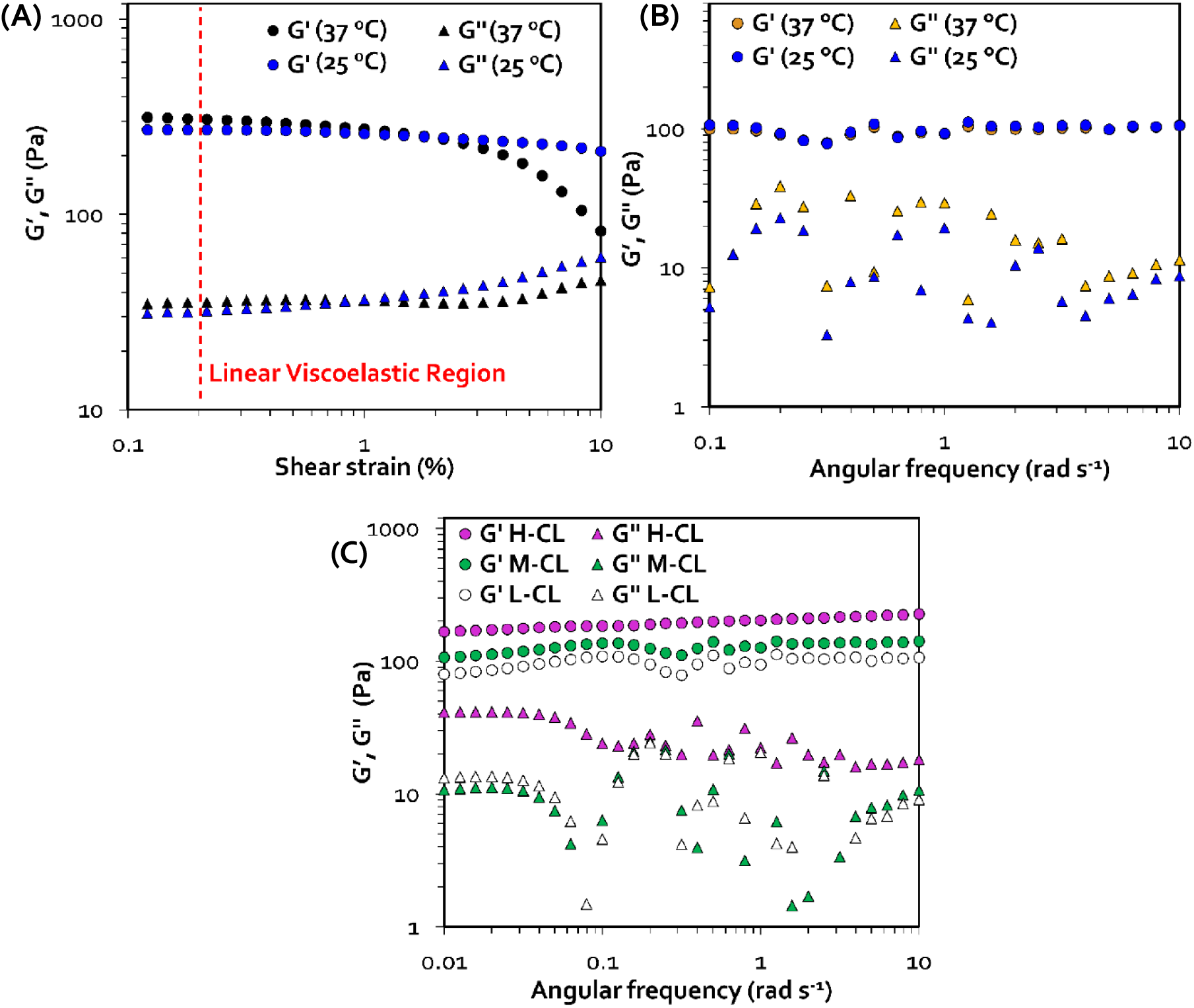
Rheological characterization of crosslinked porcine gastric mucin hydrogels. (A) Strain sweep measurements to determine the linear viscoelastic region of H-CL mucin hydrogels at RT and 37 °C by setting a constant angular frequency of 10 rad s^−1^. (B) Frequency sweep measurements to determine the shear modulus of H-CL mucin hydrogels at RT and 37 °C by setting a constant strain of 0.2%. (C) Angular frequency sweep measurements of L-CL, M-CL and H-CL crosslinked mucin hydrogels by setting a constant strain of 0.2%.

The elasticity of the hydrogels was maintained over the strain range analyzed, demonstrating the typical rheological behavior of physically and covalently-crosslinked gel-like materials that dissipate little elastic energy when sheared by virtue of their strong covalent bonds. It can be observed in **Figure 3A** that samples measured at 37 °C were less resistant to deformation at shear strains >1%, as apparent from their shorter LVE range compared to that of samples measured at RT. Following the strain sweep, a frequency sweep was performed by applying a constant strain of 0.2% to investigate the shear modulus of the three hydrogels. We initially measured the rheological properties of the hydrogels at RT and 37 °C to assess whether we can conduct the particle penetration study at RT and simplify the experimental setup. As shown in **Figure 3B**, no significant differences in the rheological properties of the hydrogels were recorded at the two different temperatures. Furthermore, the results of the frequency sweep confirmed that these hydrogels exhibit solid-like elastic properties [59], as expressed by values of storage modulus (G’) that were always ~10-fold greater than those of loss modulus (G’’) in all samples (**Figure 3B,C**). As expected, the higher the crosslinking density, the higher the G’ value and the smaller the difference between G’ and G” [60,61]. Remarkably, our hydrogels showed G’ between 70 and 200 Pa and G” between 1-40 Pa. These rheological properties resemble those of freshly extracted porcine mucin at pH values between 1 and 7 that displayed G’ and G” in the 10-200 and 2-20 Pa range, respectively [62]. These results point out that these hydrogels can serve as a reliable *in vitro* model to investigate the interaction of particulate matter with gastrointestinal mucus which is the most popular mucosal drug delivery route.

Next, we calculated the equilibrium mass-swelling ratio (Q_M_) as the weight ratio between the swollen and dry samples by gravimetric analysis and the density of the dry samples using a He pycnometer. Gravimetric measurements of the mass-swelling ratio of the different hydrogels showed that regardless of the crosslinking density, the water-content is in the range of the mucus in different mucosal tissues [13]. In addition, a slight decrease in the overall water content with increasing hydrogel crosslinking densities (and the measured G’) due to the difference in the ability of the hydrogels to retain water following the changes in the network structure and the density of the pores was observed (**Table 1**). As expected, the density of dry H-CL hydrogels was the highest of all the samples.

**Table 1.**
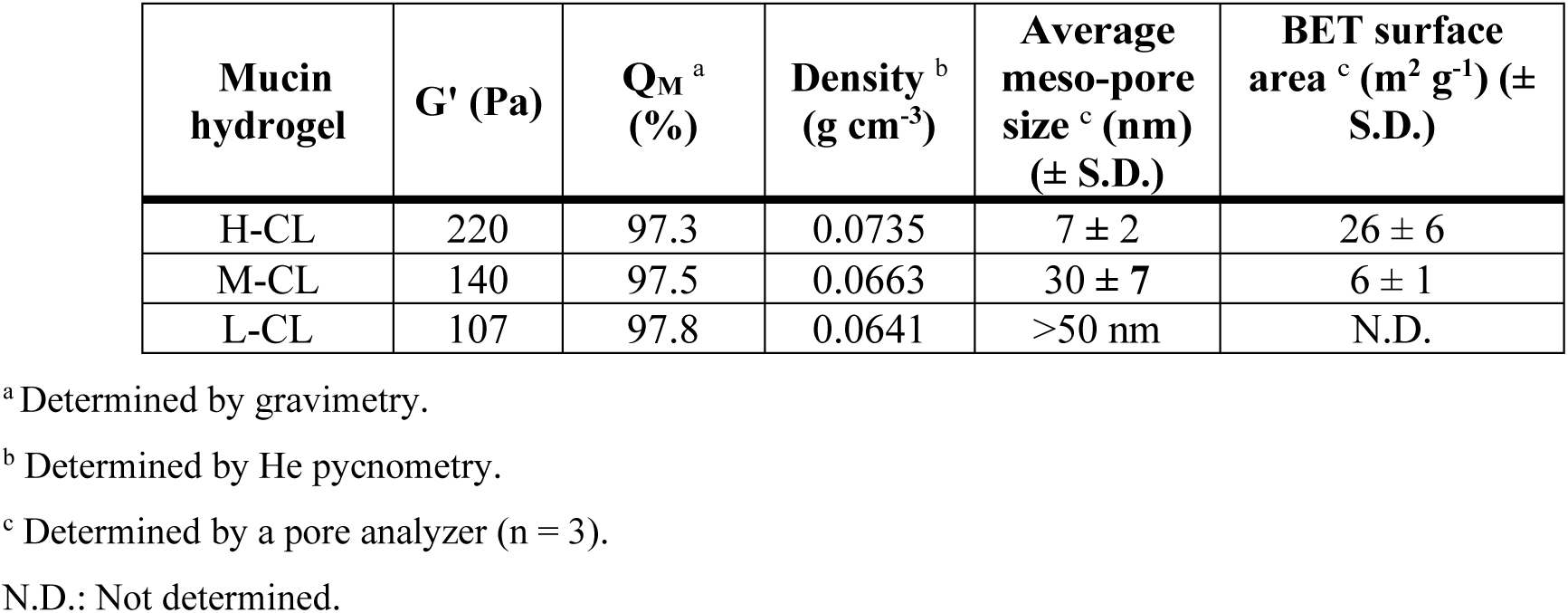
Gravimetric swelling test, He pycnometry measurements and porosimetry analysis of crosslinked mucin hydrogels. The final Irgacure^®^ 2959 concentration was 10 mg mL^−1^.

As described above, mucus is characterized by a hierarchical porous structure. To compare the structure of these synthetic hydrogels with the native one described in the literature [62], we characterized the size of the meso-pores (2-50 nm) for mucin hydrogels of different crosslinking density using a pore analyzer based on the Brunauer–Emmett–Teller (BET) theory. While the size of the pores that can be measured using this method is one order of magnitude smaller than the size of nanoparticles used in our penetration studies (see below), the goal was to obtain more structural information about the differences between the hydrogels and compare them to the features of freshly extracted gastrointestinal mucus [62]. H-CL mucin hydrogels showed an average meso-pore size of 7 nm and BET surface area of 26 m^2^ g^−1^, while M-CL counterparts of 30 nm and 6 m^2^ g^−1^ (**Table 1**). Conversely, the average pore size of the L-CL mucin hydrogel was >50 nm and could not be measured. The average pore size obtained for H-CL mucin hydrogels was in good agreement with the lower limit of pore size range in mucus of the GIT which is in the range of 10-200 nm [44,63], emphasizing once more that our hydrogels can serve as a reliable *in vitro* model of intestinal mucus. It is also worth stressing that these meso-porosity values do not rule out the presence of substantially larger pores (see below). The morphology of freeze-dried mucin hydrogels was visualized by high resolution-scanning electron microscopy (HR-SEM), as exemplified for H-CL in **Figure 4**. As expected, micrographs revealed a three-dimensional hierarchical porous structure with interconnected pores of different sizes of up to several tens of microns and shapes (**Figure 4A**), in good agreement with the literature and in analogy with the properties of native mucus [6,10,62]. M-CL and L-CL hydrogels exhibited a similar porous structure though with gradually thinner pore walls owing to a lower crosslinking density (data not shown).

**Figure 4.**
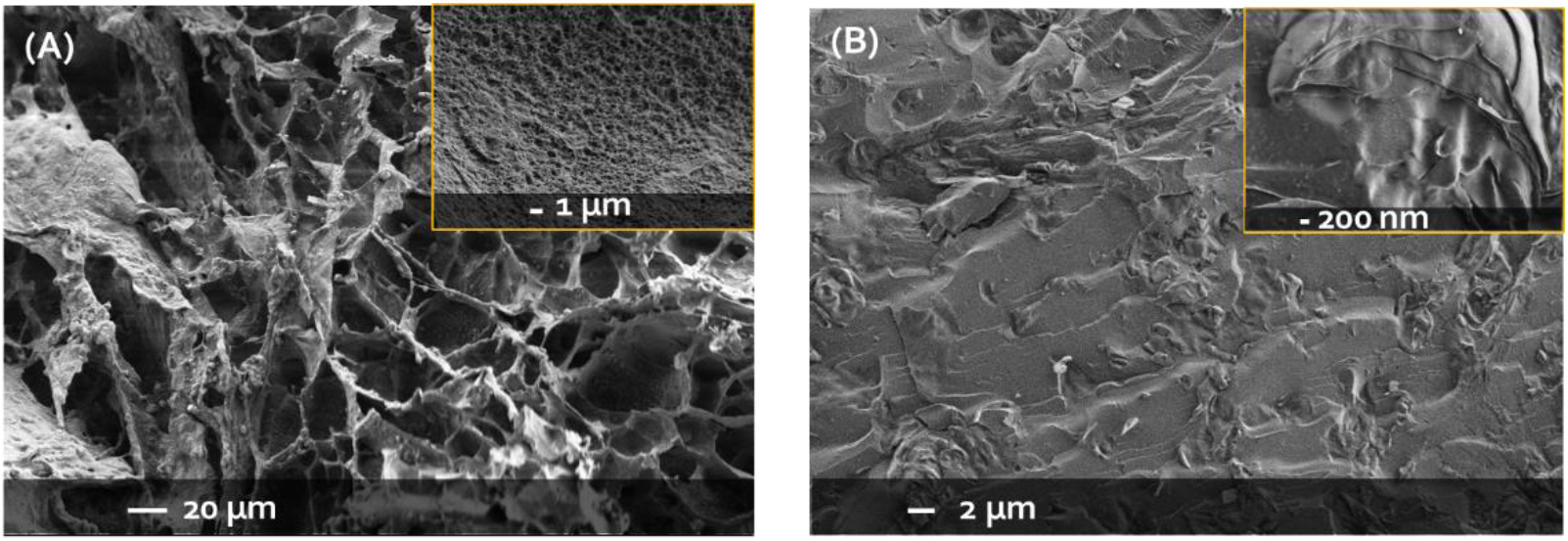
Microstructural characterization of H-CL mucin hydrogels. (A) HR-SEM micrograph of a freeze-dried hydrogel at an acceleration voltage of 1.3 kV. (B) Cryo-SEM micrograph of a swollen hydrogel at an acceleration voltage of 0.8 kV.

Since the freeze-drying of swollen hydrogels can affect the structure of the pores, we also conducted cryogenic (cryo)-SEM to visualize H-CL hydrogels in their native swollen state [64,65]. For this, samples were imaged without etching or coating at low acceleration voltage to minimally affect the original structure and network properties. A characteristic smooth inner-surface with a terraced heterogenous structure typical of biological hydrogels and the unperturbed structure of the hydrogel network were observed (**Figure 4B**) [64].

#### 2.2.2. Production and characterization of curcumin particles

For penetration studies *in vitro* we used pure curcumin particles. The rationale behind their use relied on its very low water solubility (<1 μg mL^−1^) [66] that enables the production of pure nanoparticles by nanoprecipitation in water and the ability to quantify them by fluorescence spectrophotometry without interferences due to the leaching of components of the crosslinked mucin hydrogels or photo-initiator traces into the water supernatant that could affect the quantification by UV spectrophotometry. For this, we used curcumin particles of three different sizes (expressed as hydrodynamic diameter, Dh), as measured by dynamic light scattering (DLS): 200 nm, and 1.2 and 1.33 μm. Surfactant-free pure curcumin nanoparticles were produced using a Y-shaped Si-made microfluidic device by setting flow rates of 0.2 and 2.0 mL and a volume ratio of 1:10 for the solvent and anti-solvent (water), respectively [67]. Immediately after production, the D_*h*_ (by intensity) of the nanoparticles was 200 ± 14 nm, with very small polydispersity index (PDI, a measure of the size distribution) of 0.09 and a monomodal size distribution (only one size population) (**Table S1, Figure S1A**). The physical stability of the free nanoparticles was assessed by tracking the size and the PDI over time by DLS; an increase in the measured D_*h*_ with time indicates particle agglomeration and growth which might affect the penetration studies *in vitro*. Both the D_*h*_ and the PDI remained almost unchanged for over at least six days (**Table S1**), indicating that the nanoparticles are physically stable and they can be used for penetration studies without jeopardizing their properties, which might affect the results.

To obtain larger curcumin particles, we used unprocessed curcumin microparticles with an average size of 1.2 μm, and unprocessed curcumin microparticles that were filtered to isolate a population of larger particles with an average size of 1.33 μm, as measured by DLS.

The morphological, crystallographic and thermal properties of pure curcumin particles were analyzed by HR-SEM, powder X-ray diffraction (PXRD) and differential scanning calorimetry (DSC), respectively. HR-SEM of pristine and nanonized curcumin showed irregular micron-sized structures and spherical nanoparticles, respectively (**Figure S1B,C**). In addition, pristine curcumin particles (1.2 and 1.3 μm) showed a crystalline structure with characteristic peaks that appeared at diffraction angles of 2θ equal to 7.88°, 8.90°, 12.25°, 14.58°, 17.36° and 19.49° (**Figure S1D**), in good agreement with the literature [68]. Upon nanonization, all the characteristic diffraction peaks of crystalline curcumin disappeared, indicating its amorphization (**Figure S1D**). In DSC analysis, pristine curcumin showed a melting temperature (T_m_) at 176 °C and nanonization led to a very slight shift of the T_m_ to 175 °C, and to a significant decrease of the melting enthalpy (Δ_h_) from 96.7 to 18.3 J g^−1^, confirming the increase in its amorphousness (**Figure S1E**).

#### 2.2.3. Experimental setup for particle penetration studies in vitro

After a comprehensive characterization, H-CL mucin hydrogels were used to assess the diffusion-driven particle penetration for which fresh pure curcumin particles of different size (D_h_ of 200 nm, and 1.2 and 1.33 μm) and concentration (18, 35 and 71 μg mL^−1^) were evenly dispersed on top of each hydrogel (**Figure 5A**), water poured onto the hydrogel to produce a 3 mm-height water column and the system incubated at RT; the rheological properties of mucin hydrogels at 25 and 37°C were identical, as shown above. Particle-free hydrogels subjected to an identical water column were used as control and processed following the same protocol.

**Figure 5.**
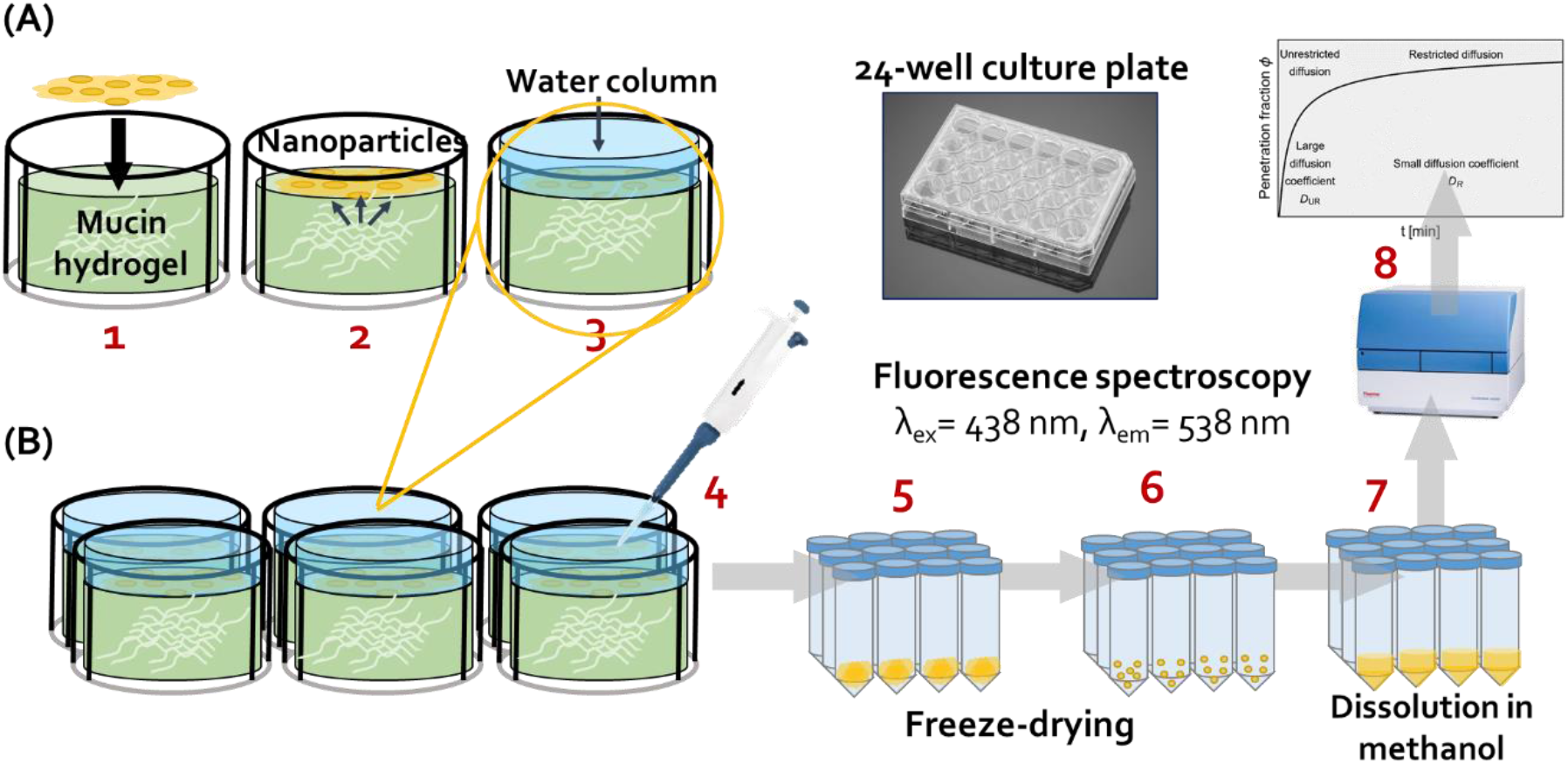
Experimental setup to assess particle penetration into mucin hydrogels that mimic the intestinal mucus. (A) Scheme of the mucin hydrogel layer inside the well plates before and after curcumin nanosuspension spreading and the application of the water column.

Particles were evenly dispersed on top of each hydrogel (1,2), and water poured onto the hydrogel to produce a 3-mm column (3). (B) Illustration of the collection of the upper water phase containing the particles that remained outside the hydrogel for freeze-drying and further analysis in the spectrofluorometer. At different time points, the supernatant above the hydrogel was carefully drawn-out and the hydrogel surface washed to ensure the recovery of 100% of the particles that did not penetrate the hydrogel (4), water aliquots containing curcumin particles were frozen in liquid nitrogen (5), freeze-dried (6), re-dissolved in methanol (7) and the concentration measured by fluorescence spectrophotometry (8).

After 5, 10, 15, 20, 40, and 60 min, the supernatant above the hydrogel was carefully drawn-out and the hydrogel surface washed to ensure the recovery of 100% of the particles that did not penetrate the hydrogel. Water aliquots containing curcumin particles were frozen in liquid nitrogen, freeze-dried, curcumin re-dissolved in methanol and the concentration measured by fluorescence spectrophotometry at an excitation wavelength (*λ_ex_*) of 438 nm and emission wavelength (*λ_em_*) of 538 nm to indirectly quantify the percentage of curcumin (as particles) that penetrated the hydrogel by calculating the difference (**Figure 5B**). The choice of H-CL hydrogels for these experiments stems from rheological properties that fit those described for native gastrointestinal mucus, and better manipulability and mechanical stability during the experiments. To investigate the effect of the crosslinking density of the hydrogel on particle penetration, we used pure curcumin nanoparticles with a D_h_ of 200 nm and at a concentration of 35 μg mL^−1^ and L-CL, M-CL and H-CL hydrogels. In each set of experiments, only one parameter, namely particle size, particle concentration, or crosslinking density of the mucin hydrogel, was changed while keeping the others fixed to study the effect of each parameter separately.

#### 2.2.4. Prediction model for particle penetration via diffusion into hydrogels

To study the influence of key factors on the overall diffusion of particles into hydrogels, we considered a purely diffusion-driven process, where no external forces are applied. We focus on three parameters: (i) the particle size, which is quantified by the average hydrodynamic diameter of a particle (D_h_), (ii) the concentration of particles in the medium (C), and (iii) the crosslinking density of the hydrogel, which is described by the shear storage modulus (G’). Recall that in gels, the relation between the shear modulus and the crosslinking density *N*_0_ is given by is given by 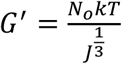 [69–71]. In physically crosslinked biological networks, crosslinks can dissociate but the stiffness still exhibits a dependency on the crosslinking density [72,73]. Based on preliminary experiments, the rheological properties of intestinal mucus, the average size of nanonized drugs (<1 μm), and the dose of orally administered drugs, we defined the characteristics of a reference system as follows: D_h_ = 200 nm, C_R_ = 35 μg mL^−1^, G’ = 220 Pa, and ϕ_0,R_ = 55%.

To determine the overall sensitivity *γ*_•_, we modify the parameter • = D_h_, C or G’ while keeping the others fixed and perform a fit using least squares. Recall that *α* is a constant that characterizes the effective diffusion coefficient. It is emphasized that while the use of Eq. 2 is simple, its limitation is that the value of *γ*_•_ depends on the reference system.

##### Effect of particle size

Size filtering is one of the predominant factors that dictate particle penetration *via* diffusion into biological gels [44,74]. In the first set of experiments, we focused on the ability of the mucin hydrogels to hinder the penetration of particles based on their size. For this, we used three different curcumin particles sizes and the fraction of particles that penetrated into the hydrogel was assessed by measuring the fluorescence of curcumin particles that remained in the water supernatant (that did not penetrate), as described in **Figure 5B**. The experimental results and their fit are presented in **Figure 6A**. The fitting parameter and the computed value of α were γ_D_h__ = 0.5 μm^−1^ = 0.5 and 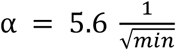, respectively. The size distribution and populations (in percentage) of each group, as measured by DLS, are presented in **Figure 6B**.

**Figure 6.**
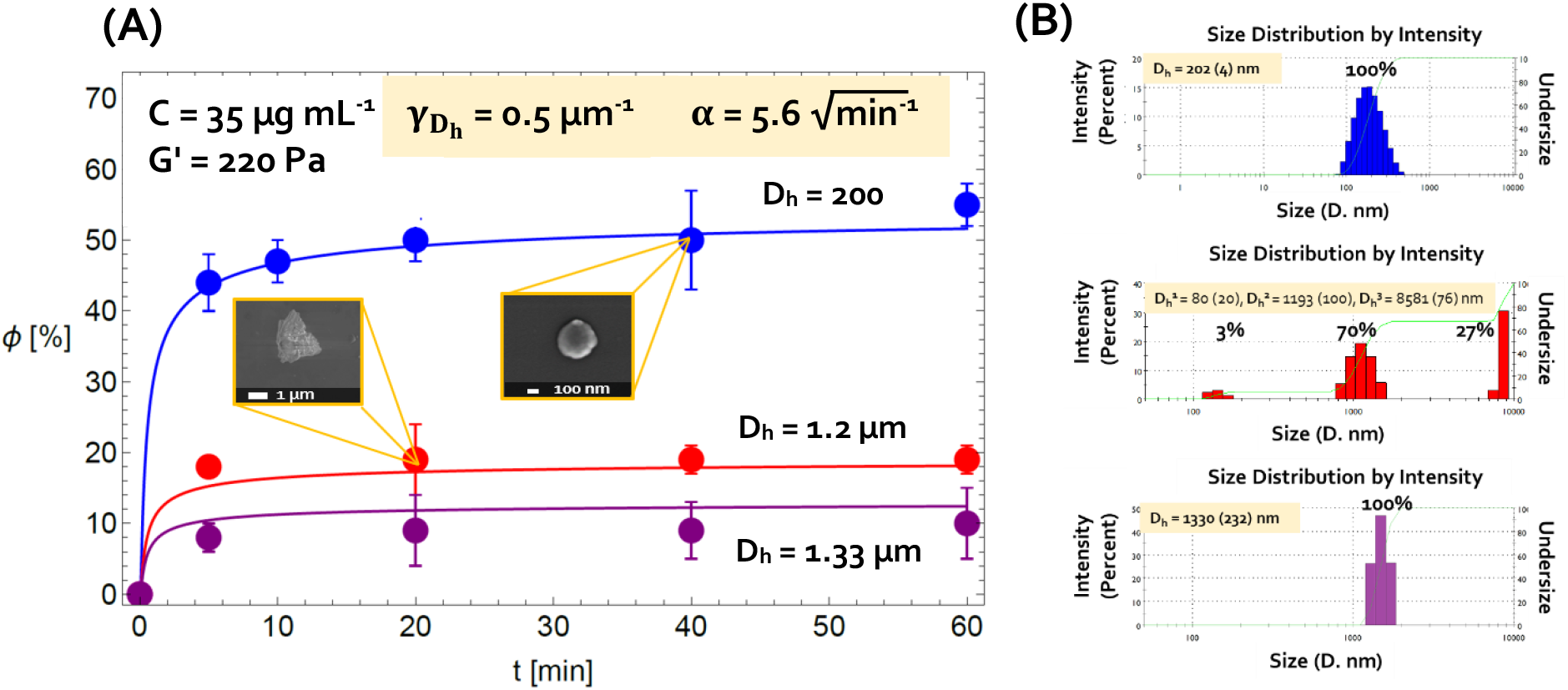
Effect of particle size on particle penetration into H-CL mucin hydrogels. (A) Experimental results and modeling of the fraction of curcumin particles that penetrated the hydrogel (in percentage) for three representative average particle sizes, D_h_. (B) Size distribution and populations, as measured by DLS.

Differences based on size were significant, as demonstrated by the sharp change in the fraction of particles that penetrated the H-CL mucin hydrogel over time. For example, after 5 min the fraction of curcumin nanoparticles that penetrated into the hydrogel was 44%, whereas only 18% and 8% of the particles with sizes of 1.2 and 1.33 μm, respectively, did so. Later on, the percentage of particles that penetrated into the hydrogel remained almost constant even for 200-nm nanoparticles, strongly suggesting that once part of the penetration sites in the hydrogel are filled with particles, further particle penetration is precluded. In addition, the time required for this phenomenon to occur is very short (5 min) compared to the average intestinal transit time and that time is not the limiting stage of the penetration process. These observations were in good agreement with previous reports for the diffusion of particles larger than 500 nm through reconstituted gastrointestinal porcine mucus *ex vivo* showing that although the diffusion of these particles was limited, it was not completely prevented [44,74], strengthening our observations of particles larger than 1 μm that can penetrate mucus-mimicking mucin hydrogels to some extent. These results demonstrate that the change in particle dimensions can significantly increase/decrease the transport rate of the particles across mucus and highlight the relevance of fine-tuning the size of PDNPs for oral administration.

##### Effect of particle concentration

Another physicochemical parameter that is less frequently investigated and could have a striking effect on the penetration of particles into mucus is particle concentration (which in preclinical and clinical studies depends on the drug dose). To investigate this phenomenon, we studied the response with three concentrations of pure curcumin nanoparticles (D_h_ = 200 nm), namely C = 18, 35, and 71 μg mL^−1^ that corresponded to nanoparticle nanoparticle concentrations per hydrogel surface area of 5.2, 10 and 20 μg cm^−2^, respectively. As shown in the preliminary oral pharmacokinetics (PK) experiments, these concentrations per surface area fit those obtained after the oral administration of clinically relevant doses (see below). The experimental results are illustrated in **Figure 7** and the fitting parameter γ_C_ =5.7 · 10^−3^ mg mL^−1^.

**Figure 7.**
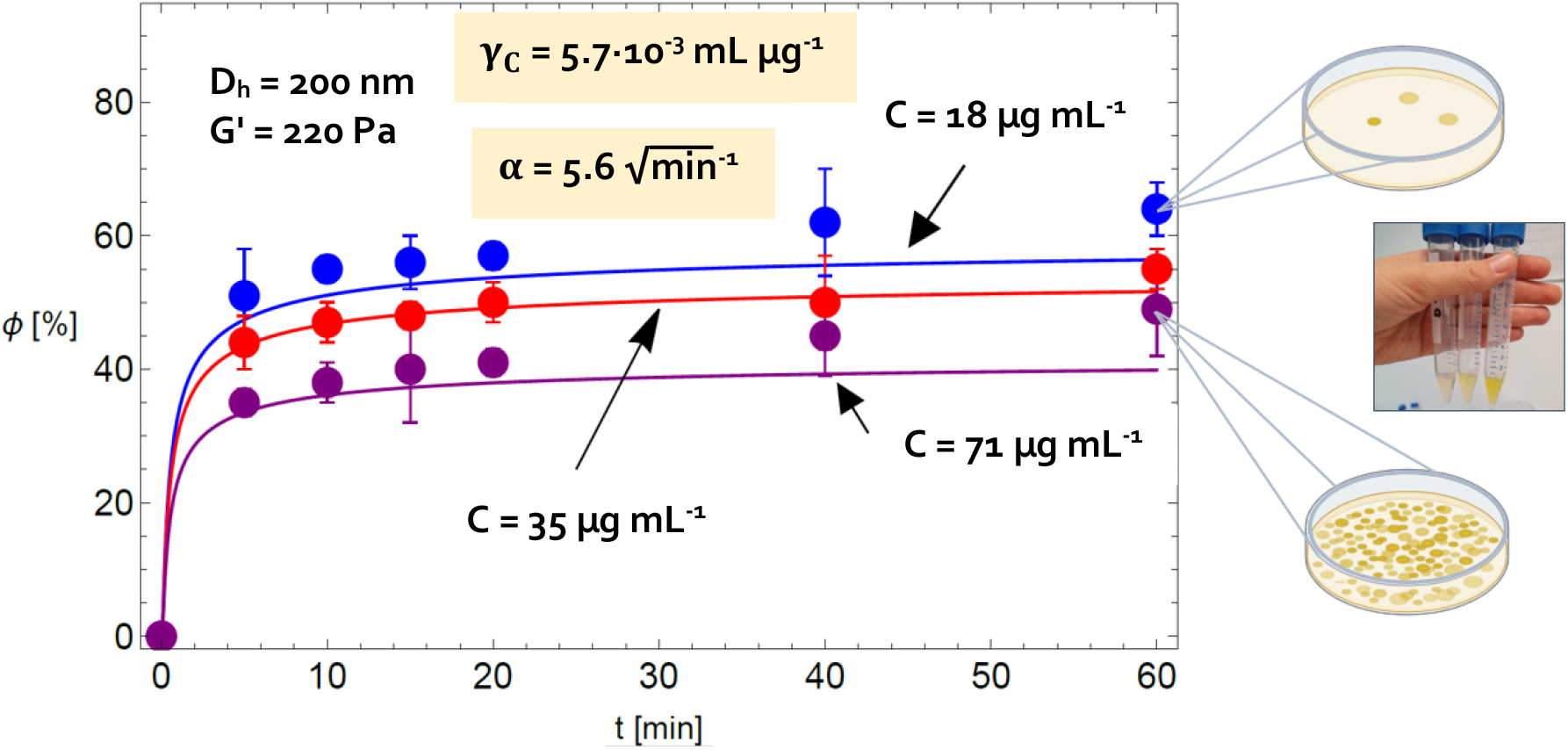
Effect of particle concentration on particle penetration into H-CL mucin hydrogels. Experimental results and modeling of the fraction of curcumin particles that penetrate the hydrogel (in percentage) for the three representative concentrations C.

Interestingly, **Figure 7** shows that the percentage of particle penetration decreases as the concentration C increases. This counterintuitive behavior stems from the local particle-particle and particle-hydrogel interactions. To understand this, we highlight that the overall number of particles that simultaneously attempt to penetrate the hydrogel increases with the concentration. Consequently, neighboring particles (or particle-particle interactions) interrupt the diffusion process in two ways: (i) “exhaustion” of penetration sites due to saturation of particles along the hydrogel surface and (ii) increase in the local intermolecular spacing due to local deformations of chains in response to particle penetration translates into a decrease in nearby spacings. This reduction prevents the penetration of additional particles. Note that the latter mechanism can hinder the penetration of neighboring particles even when the surface of the hydrogel is not completely saturated by them. It is worth noting that based on physical stability data measured by DLS, pure curcumin nanoparticles in suspension do not undergo agglomeration and size growth during the time of the assay. Thus, a decrease in particle penetration at greater concentrations was not associated with particle agglomeration.

To quantify the effect of particle concentration on the overall diffusion process, we assume that the particles are homogeneously distributed over the surface of the hydrogel. The overall number of particles in the system is 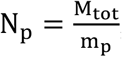 where m_p_ = ρ_p_V_p_ and M_tot_ are the mass of particle and the total mass, respectively. The density ρ_p_ = 1.3 mg mm^−3^ and V_p_ is the volume of a particle. The area fraction that particles with a diameter D_h_ occupy along the surface of the hydrogel is

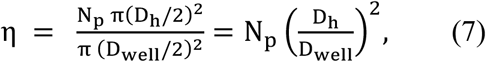

where D_well_ = 15.6 mm is the diameter of the 24 well-plate (according to the specifications of the manufacturer).

To estimate the maximum value of η (Equation 7), or the maximum area coverage that can be obtained, and the average distance between particles, we consider a representative unit with the dimensions L × L comprising one effective particle, as shown in **Figure 8**. We denote the distance between particles by Δ_p_ such that L = D_h_ +Δ_p_. Accordingly, 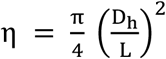. The maximum packing is obtained when the distance between the particles Δ_p_= 0 (and L = D_h_), resulting in η_max_ ≈ 78.5%. In addition, the average distance between particles is

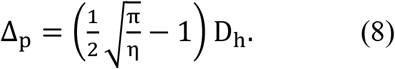

**Figure 8.**
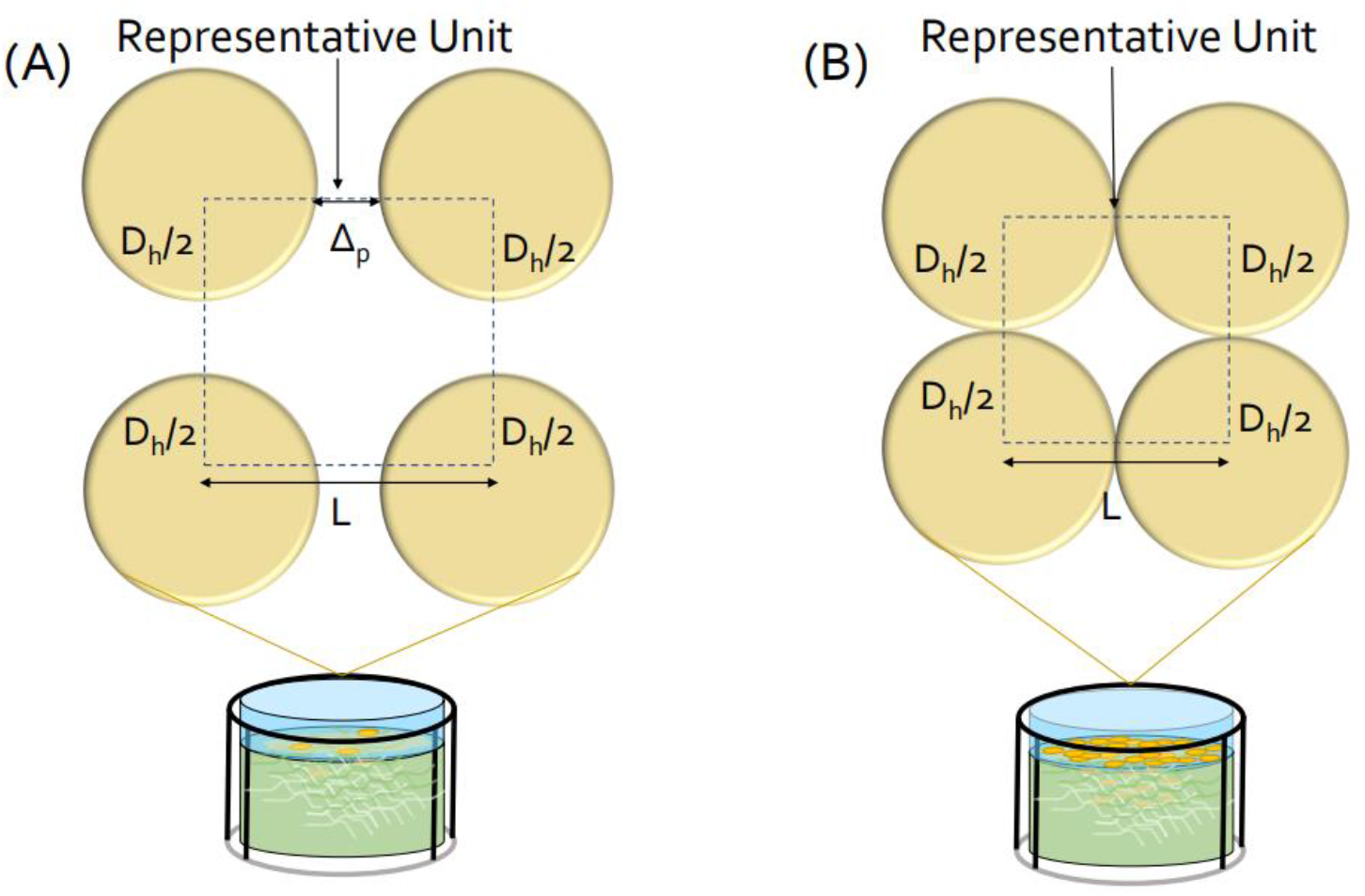
Schematic representation of the arrangement of the maximum spherical particles that can be contained inside a 2D representative structural unit during particle penetration in two different conditions. (A) The hydrogel surface is partially covered with particles (η < η_max_), and (B) full particle coverage and saturation of the hydrogel surface layer (η > η_max_).

The three concentrations considered in **Figure 7** correspond to a particle with a diameter of D_h_ = 200 nm and the overall particle masses M_tot_ = 0.01,0.02, and 0.04 mg. Using Equation 7, we obtain surface area coverages of η_18_ = 30.2%, η_35_ = 60.4%, and η_71_ = 120.7% for particle concentrations 18, 35, and 71 μg mL^−1^, respectively. The value of and η_71_ exceeds the maximum possible packing, implying that for a concentration of 71 μg mL^−1^, the hydrogel surface is packed with layers of particles. As a consequence, all of the particles interact with each other and reduce the overall penetration, as experimentally observed in **Figure 7**. Saturation of the hydrogel surface is expected to affect the *in vivo* performance of the nanoparticles for growing concentrations (doses).

By employing Equation 8, we find that the average distance between particles for the concentrations 18 and 35 μg mL^−1^ is approximated as such that Δ_P_18__ ≈ 122 nm and Δ_P_35__ = 28 nm, respectively. Since in both cases Δ_p_ < D_h_, it is clear that there is some influence of the particles on the overall diffusion process.

The above analysis emphasizes that the interaction between the particles in the hydrogel plays a role in the diffusion process and the overall percentage of penetrated particles. Specifically, these interactions interfere the diffusion by locally deforming chains and decreasing interchain spacings.

##### Effect of hydrogel crosslinking density

Mucus displays a different set of physicochemical and rheological properties based on the mucosal tissue and portion [75]. The crosslinking density of biological gels such as mucus governs their structural and mechanical properties and can be easily tuned *in vitro* by changing the crosslinker concentration and/or by changing the conditions of the reaction [57,76,77].

In the photo-initiated crosslinking of the mucin-MA to produce hydrogels, the photo-initiator concentration and the relative volume of the solvent in which it is dissolved can be changed to control the crosslinking density of the hydrogels [57]. To produce H-CL, M-CL and L-CL mucin hydrogels, we kept the final concentration of the photo-initiator in the precursor mucin-MA aqueous solution constant and changed the volume of photo-initiator ethanolic solution added so the final ethanol concentration in this solution changes. Then, we assessed the penetration of pure curcumin nanoparticles (D_h_ = 200 nm) at a concnetration of 35 μg mL^−1^ into hydrogels with variable crosslinking density. As anticipated, an increase in the crosslinking density leads to a significant reduction in the fraction of nanoparticles that penetrate the hydrogel over time (**Figure 9**). This is due to the reduction in the intermolecular spacing with increasing crosslinking density, which restricts the deformation range of the polymer chains. The more crosslinked the structure of the network (and the greater the number of crosslink sites), the denser it becomes, and thus the average interchain spacing (or effective pore-size) between crosslinks becomes smaller. Consequently, it is harder for the diffusing particles to stretch the chains and penetrate the network. The calculated fitting parameter which demonstrates the overall response of the particle-hydrogel system to the change in the crosslinking density of the hydrogel for this set of experiments was γ_G’_ = 4.4 · 10^−3^.

**Figure 9.**
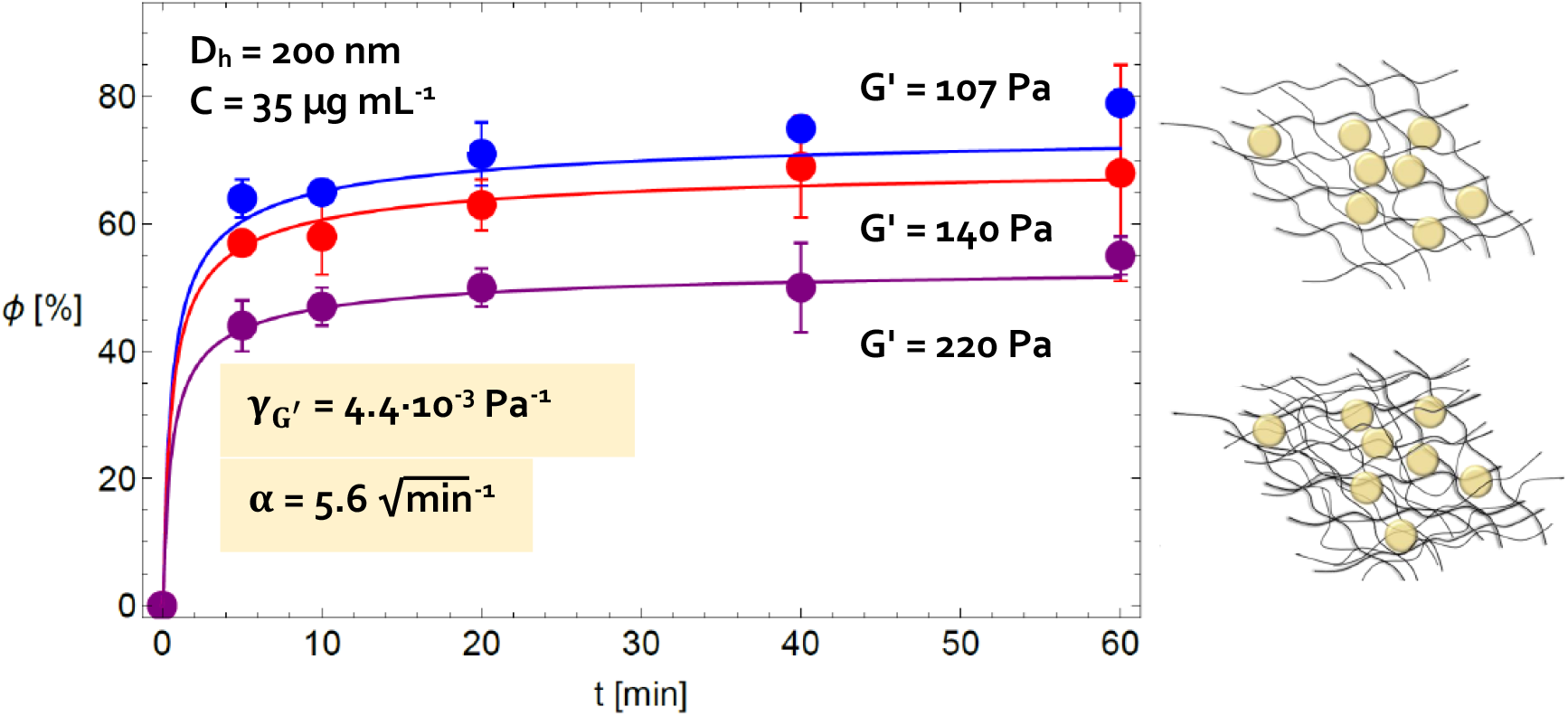
Effect of the crosslinking density of mucin hydrogels on particle penetration. Experimental results and modeling of the fraction of curcumin nanoparticles (200 nm) that penetrate the hydrogel (in percentage) for three representative crosslinking densities, which are characterized by three values of the shear storage modulus, G’, as measured in rheology experiments.

#### 2.2.5. Visualization of pure curcumin nanoparticles in the mucin hydrogels

After assessing the effect of fundamental parameters on particle penetration into mucus-mimicking mucin hydrogels, we visualized the hydrogels 30 min after exposure to pure curcumin nanoparticles (D_h_ = 200 nm) at a concnetration of 35 μg mL^−1^ by cryo-SEM and evidenced the presence of newly formed cavities (**Figure 10A**, see yellow arrows) which indicates the existence of curcumin nanoparticles within the hydrogel matrix (**Figure 10B**).

**Figure 10.**
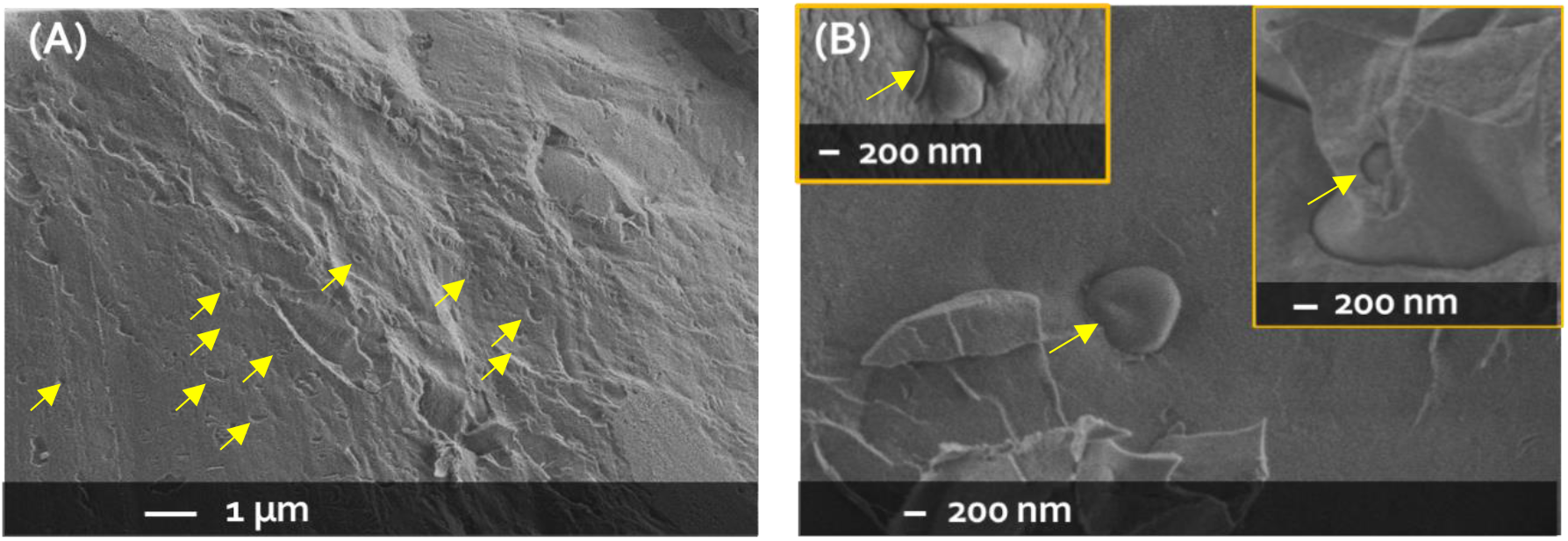
Characterization of H-CL mucin hydrogels upon exposure to pure curcumin nanoparticles in penetration experiments by cryo-SEM. (A) Structure of the mucin hydrogel showing the cavities left by the nanoparticles that penetrated it (see yellow arrows). (B) Pure curcumin nanoparticles that penetrated into the mucin hydrogel (see yellow arrows). Different regions of the hydrogel are shown. Low acceleration voltage of 0.8 kV was used. The nanoparticles showed a D_h_ of 200 nm and the concentration was 35 μg mL^−1^.

### 2.3. Oral pharmacokinetics of a model drug in rats

An advantage of developing experimental-theoretical tools for the prediction of the interaction between particles and hydrogels based on fundamental features such as particle size and concentration, and the crosslinking density of the network is that they could be applied regardless of the chemical nature of the particle and thus, in the field of pharmaceutical research and development, for different drugs. After deriving the physical models and correlating the theory with the experimental results from the particle penetration assays *in vitro*, we challenged our experimental-theoretical approach *in vivo*. For this, we conducted a preliminary study of the oral PK in rat of a clinically relevant poorly water-soluble tyrosine kinase inhibitor of oral administration, dasatinib free base (DAS), used in the therapy of chronic myeloid and acute lymphoblastic leukemias and in autoimmune diseases in patients with resistance or intolerance to prior therapy [78,79]. The oral bioavailability of DAS is in the 14-34% range [80], which justifies its nanonization to increase it.

Since the properties of intestinal mucus cannot be tuned *in vivo*, we assessed the effect of particle size and concentration. To study the effect of particle size, we administered (i) DAS nanoparticles with D_h_ of 200 nm and (ii) unprocessed DAS particles with a D_h_ of 1.2 μm (similar to pure curcumin nanoparticles of intermediate size), both at a dose of 10 mg kg^−1^, which is clinically relevant [81,82]. To study the effect of particle concentration, we administered two different doses (10 and 15 mg kg^−1^) of pure DAS nanoparticles (D_h_ = 200 nm). The rats used in our studies weighed 230 ± 34 g and thus, for clinically relevant doses of 10 mg and 15 mg kg^−1^, the average amount of orally administered DAS was 2.30 ± 0.34 and 3.45 ± 0.51 mg, respectively. Intestinal surface area is a key parameter in the absorption rate of drugs. Meshkinpour *et al*. calculated the mean surface area of the intestine of rats experimentally and showed that it increases linearly during the first 6 weeks of life and it reaches a plateau of 142 cm^2^ when the average weight is 177 ± 8 g [83]. A further weight increase does not change the mean intestinal surface area. According to this, the estimated DAS particle concentration per surface area for doses of 10 and 15 mg kg^−1^ was 16 ± 2 and 24 ± 3 μg cm^−2^, respectively, these values being similar to the concentration per surface area of pure curcumin nanoparticles with C of 35 and 71 μg mL^−1^ (that corresponded to 10 and 20 μg cm^−2^, respectively) and that were used in the particle penetration assay *in vitro*.

Pure DAS nanoparticles were produced using the same microfluidics method described for curcumin and their size and morphology were very similar to those of curcumin counterparts, as reported elsewhere [67]. In addition, pure curcumin and DAS nanoparticles showed a zeta-potential of −18 mV (**Table S2, Figure S2**), so the effect of the surface charge was neglected.

Plasma concentration–time plots and PK parameters following oral administration are presented in **Figure 11** and **Table 2**, respectively. The plasma DAS concentration upon administration of unprocessed drug (dose of 10 mg kg^−1^) increased quickly, the C_max_ was 72.5 ng mL^−1^ and the time to the C_max_ (t_max_) 4.0 h. Then, the drug was cleared fast with a t_1/2_ of 5.1 h (**Figure 11**, **Table 2**). These results were in good agreement with the literature [80,84].

**Figure 11.**
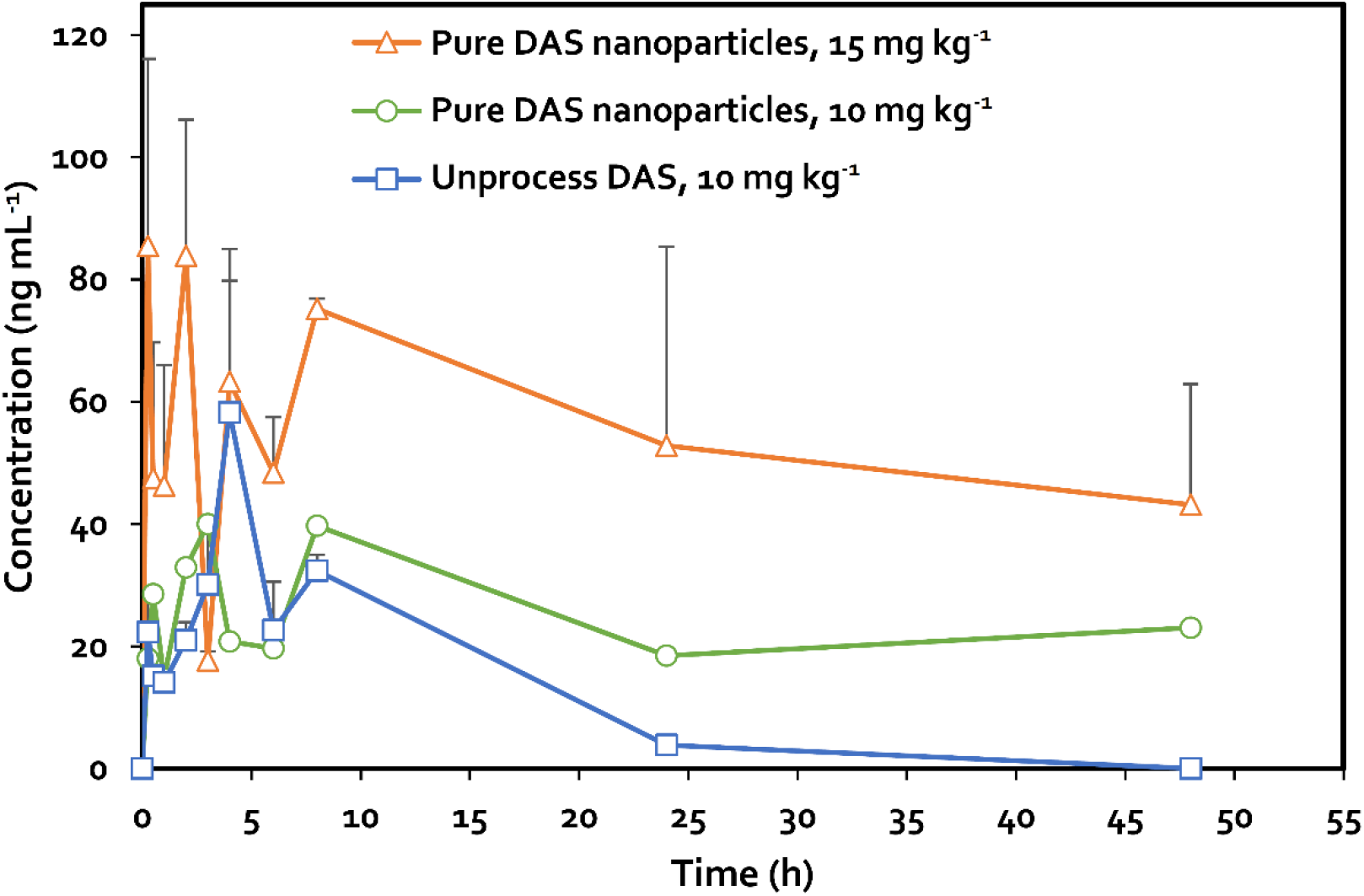
Oral pharmacokinetics of dasatinib (DAS) in Sprague-Dawley rats. Mean plasma concentration versus time profiles following the oral administration of (A) one single dose (10 mg kg^−1^) of (■) unprocessed DAS (D_h_ = 1.2 μm) and (•) pure DAS nanoparticles (D_h_ = 200 nm) to rats (n = 6). (B) Comparative pharmacokinetics of pure DAS nanoparticles (D_h_ = 200 nm) in two different doses (•) 10 mg kg^−1^and (▲) 15 mg kg^−1^. Results are expressed as mean ± S.E.

**Table 2.** Oral pharmacokinetics of dasatinib (DAS) in Sprague-Dawley rats. Pharmacokinetics parameters after oral administration of a single dose of unprocessed DAS (■) and DAS PDNPs to Sprague-Dawley rats (n = 3).

| PK Parameter | Raw DAS $10 \text{ mg kg}^{-1}$ | | PDNPs $10 \text{ mg kg}^{-1}$ | | PDNPs $15 \text{ mg kg}^{-1}$ | |
| --- | --- | --- | --- | --- | --- | --- |
|  | Mean | CV (%) | Mean | CV (%) | Mean | CV (%) |
| $C_{\text{max}}$ ($\text{ng mL}^{-1}$) | 72.5 | 27 | 40.0 | 15 | 85.5<sup>b</sup> | 27 |
| $AUC_{0-\infty}$ ($\text{ng h mL}^{-1}$) | 604.8 | 9 | 4714.9<sup>a</sup> | 46 | 5824.5 | 47 |
| $t_{\text{max}}$ (h) | 4.0 | 0 | 3.0 | 122 | 0.25 | 97 |
| $t_{1/2}$ (h) | 5.1 | 40 | 106.7<sup>a</sup> | 46 | 51.5 | 47 |
| $F_r$ | 1.0 | - | 7.8 | - | - | - |

<sup>a</sup> Statistically significant difference ( $p < 0.05$ ) when compared with the $10 \text{ mg kg}^{-1}$ unprocessed drug.
<sup>b</sup> Statistically significant difference ( $p < 0.05$ ) when compared with the $10 \text{ mg kg}^{-1}$ pure drug nanoparticles.

When the same dose was administered using the nanoparticles, the C_max_ decreased to 40.0 ng mL^−1^ and the t_max_ was shortened to 3.0 h though the t_1/2_ showed a dramatic prolongation to 106.7 h (a 20.9-fold increase) (**Table 2**). Moreover, DAS nanonization resulted in a statistically significant 7.8-fold increase of the area-under-the-curve between 0 and ∞ (AUC_0-∞_) from 604.8 to 4714.9 ng h mL^−1^ with respect to the unprocessed drug and the plasma concentration remained within the 10-20 ng mL^−1^ range for at least 48 h. The effective DAS concentration in plasma required to inhibit 90% of phospho-BCR-ABL in chronic myeloid leukemia is 10.9 ng mL^−1^ in mice and 14.6 ng mL^−1^ in humans [82], indicating that the nanoparticles ensure therapeutic concentrations for a longer time. This increase in the oral bioavailability and prolongation of the t_1/2_, as previously reported for indinavir free base [24], suggest that DAS nanoparticles are retained in the intestinal mucosa and undergo slow dissolution and absorption over time. These results are in good correlation with our theoretical models and our experimental findings *in vitro*, suggesting that nanonization leads to a significant increase in the number of particles that manage to penetrate the mucin hydrogel network because the hydrogel is not saturated by them, as shown for a C = 35 μg mL^−1^ (and concentration per surface area of 10 μg cm^−2^) (**Figure 6**).

The comparison between the two different doses of pure DAS nanoparticles (10 and 15 mg kg^−1^) showed that the overall sustained release profile was conserved, and a statistically significant 2.1-fold increase in the C_max_ from 40.0 ng mL^−1^ to 85.5 ng mL^−1^ was observed. At the same time, the t_max_ and t_1/2_ were shortened to 0.25 h and 51.5 h, respectively, and the AUC_0-∞_ showed a 1.2-fold increase, which was statistically insignificant when compared to the AUC_0-∞_ of the nanoparticles at a lower dose. These findings are associated with the saturation of the nanoparticle entrapment sites in the intestinal mucus which results in a faster absorption and elimination of the unabsorbed drug with respect to the lower dose of pure DAS nanoparticles (10 mg kg^−1^). Despite the different experimental conditions of the penetration studies *in vitro* (quasi-static) and of the oral administration *in vivo* (dynamic), PK results with a dose of 15 mg kg^−1^ (and a DAS particle concentration to surface area of 24 ± 3 μg cm^−2^) are in good agreement with the behavior shown by the most concentrated sample of pure curcumin nanoparticles used in the penetration studies *in vitro* (71 μg mL^−1^) in which the saturation of the penetration sites due to the restrictions posed on neighboring particles that penetrate the hydrogel layer simultaneously was observed (**Figure 8**). To further validate our experimental-theoretical tools, similar PK studies with different poorly water-soluble drugs and doses will be conducted in the future.

## 3. Conclusions

Aiming to understand the contribution of different key particle properties to the interaction with mucosal tissues, the present work investigated the interaction between particles and an *in vitro* model of intestinal by utilizing both experimental and theoretical approaches. We envision the particle penetration into mucus as a spontaneous two-stage process and propose a phenomenological model that account for the physics behind this diffusion-derived process and capture the dependence of our particle-hydrogel system on changes in each one of the three key parameters assessed (particle size, particle concentration, and hydrogel crosslinking density) to complement our experimental findings. Finally, we challenged our experimental-theoretical approach *in vivo* by conducting a preliminary oral pharmacokinetic study with a clinically relevant anti-cancer drug in rat. Overall, these results reflected well the *in vitro* findings and complemented our derived theoretical models, revealing the potential of this work to shed light on the interaction between nanomaterials and mucosal tissues. We envision that the insights of this work will pave the way for the understanding and the prediction of the interactions between different types of nanoparticles and biological gels such as mucin based on the criteria assessed and will set the foundation for a more rational approach for the design of nanomedicines for mucosal drug delivery as well as the assessment of risk factors in the field of nano-safety.

## 4. Experimental section

### 4.1. Synthesis and characterization of mucin hydrogels

#### 4.1.1. Mucin methacrylation

Mucin hydrogels of different crosslinking density were prepared by using a synthesis previously reported for bovine submaxillary mucin with modifications [46]. To form a covalently crosslinked mucin hydrogel, porcine gastric mucin type II (Lot number M2378, Sigma–Aldrich, St. Louis, MO, USA) was methacrylated with MA (Sigma–Aldrich) to obtain mucin-MA. For this, 2 g of mucin was dissolved in water to obtain a final concentration of 10 mg mL^−1^. The pH of the solution was adjusted to 8.0 with a 10 N NaOH (Bio-Lab Ltd., Jerusalem, Israel) solution in water. Then, the dissolved mucin was cooled on ice for 30 min (until its temperature reached 5-8 °C) before the addition of MA at a mucin:MA weight ratio of 1:2. Immediately after adding the MA, the pH of the solution was again adjusted to 8.0 to ensure optimal reaction conditions [46]. Then, the solution was gently stirred overnight at 4 °C. The day after, the solution was centrifuged (10 min, 750 RPM, SW28 rotor, Laborgeräte Beranek GmbH, Weinheim, Germany) to remove excess MA and mucin precipitates and dialyzed against distilled water in regenerated cellulose dialysis membranes (molecular weight cut-off - MWCO - of 12-14 kDa, Spectra/Por^®^ 4 nominal flat width of 75 mm, diameter of 45 mm and volume/length ratio of 18 mL cm^−1^, Spectrum Laboratories, Inc., Rancho Dominguez, CA, USA) for 2 days with two water changes per day. The resulting mucin-MA was immediately frozen in liquid nitrogen and freeze-dried (Labconco Free Zone 4.5 plus L Benchtop Freeze Dry System, Labconco, Kansas City, MO, USA) and stored at 4 °C until use.

To quantify the degree of mucin methacrylation of mucin-MA, the TNBSA assay was performed. Mucin methacrylation consumes the primary amine groups on the glycoprotein backbone, leaving less amine groups available for the reaction with TNBSA. For this, freeze-dried mucin-MA and unreacted mucin samples were separately dissolved at 1 mg mL^−1^ in 0.1 M sodium bicarbonate buffer pH 8.5 (Merck, Kenilworth, NJ, USA). Then, 0.5 mL of each sample solution was mixed with 0.01% TNBSA solution (Tzamal, Petach Tikva, Israel) in 0.1M sodium bicarbonate buffer and incubated for 2 h, at 37 °C. Next, 0.25 mL of 10% w/v sodium dodecyl sulfate (Sigma-Aldrich) and 0.125 mL of 1 M hydrochloric acid (Bio-Lab Ltd.) were added to stop the reaction. The absorbance of each sample was measured using UV-Vis spectrophotometry (Multiskan GO Microplate Spectrophotometer with Skanlt™ software (Thermo Fisher Scientific Oy, Vantaa, Finland) at 335 nm and interpolated in a calibration curve of glycine (Sigma-Aldrich) in water in the 1-16 μg mL^−1^ range (R^2^ = 0.999) was plotted to determine the number of free amine groups before and after the methacrylation reaction.

#### 4.1.2. Preparation of mucin hydrogels

To prepare mucin hydrogels, freeze-dried mucin-MA was dissolved in water at a concentration of 40 mg mL^−1^ and crosslinked under UV light (365 nm, B-100AP model UV lamp 20,000 μW cm^−2^, Analytik Jena US LLC, Upland, CA, USA) for 8 min using Irgacure^®^ 2959 (Sigma– Aldrich) as photo-initiator (10 mg mL^−1^). To obtain hydrogels of different crosslinking density, porosity and rheological properties, the volume percentage of ethanol in the total volume of the mucin-MA precursor varied between 10%, 20%, and 30% *v/v* for the L-CL, M-CL, and H-CL mucin hydrogels, respectively. After UV crosslinking, fresh hydrogels were left at RT for 24 h to allow the free radical polymerization reaction to reach an equilibrium state.

#### 4.1.3. Characterization of mucin hydrogels

##### Water content

A gravimetric swelling test was performed on the mucin hydrogels of different crosslinking density to determine their water content. For this, hydrogels were soaked in water and allowed to swell to equilibrium at 37 °C for 24 h. Then, the excess of water was removed, and their wet weight (M_s_) was recorded. Samples were then freeze-dried overnight, and their weight (M_d_) recorded again. The equilibrium mass-swelling ratio (Q_M_) was calculated as the ratio of the mass of the swollen hydrogel and dry samples. Results are presented as the average of triplicates.

##### Rheological testing

To assess the rheological properties of mucin hydrogels, a frequency sweep with a constant strain and a strain sweep with a constant angular frequency was performed using a shear rheometer (MCR302 Anton-Paar Rheometer, Graz, Austria). For this, hydrogels with a diameter of 26 mm were prepared, soaked in water for 24 h before the analysis and placed between parallel steel plates of the rheometer (plate size 25 mm) for the sweep tests. Sand blasted plates were used to prevent wall slip. The shear strains and frequencies during testing ranged from 0.1-10% and 0.1-10 rad s^−1^, respectively, and the tests were performed at both RT and 37 °C.

##### Meso-porosity

The meso-porosity of the of L-CL, M-CL and H-CL mucin hydrogels was measured using pore analyzer (Micromeritics TriStar II 3020, Micromeritics Instrument Corp., Norcross, GA, USA) based on the calculation BET theory. Samples were vacuum freeze-dried before testing and test temperature was 30 °C. Results are presented as Mean ± S.D. (n = 3).

##### Density

The density of crosslinks of L-CL, M-CL and H-CL mucin hydrogels was determined using a high-precision He pycnometer (Micromeritics AccuPyc II 1340, Micromeritics Instrument Corp.). Samples were freeze-dried prior to testing and purged to exclude moisture and adsorbed impurities before conducting the measurements. Results are presented as the average of triplicates.

##### Microstructural analysis

To visualize the porous structure of the H-CL mucin hydrogels, freeze-dried samples were cut with razorblade and their cross-section was placed on top of a double-sided carbon tape and visualized by HR-SEM (acceleration voltage of 1 - 4 kV, Ultraplus, Carl Zeiss SMT GmbH, Oberkochen, Germany). Conductive silver paint (SPI#05002-AB silver conductive paint 1.0 Troy oz., SPI Supplies, West Chester, PA, USA) was applied at the corners of each sample. Mucin hydrogels in their native swollen state were visualized using a Zeiss Ultra Plus High-Resolution Cryo-SEM (Carl Zeiss SMT GmbH) equipped with a Schottky field-emission electron gun and a Leica VT100 cryo-system. The specimens were maintained at −145 °C and imaged without coating at a low acceleration voltage and under low-dose imaging conditions, to prevent specimen charging and radiation damage. To avoid freezing artifacts during specimen preparation high-pressure freezing was used to reduce the rate of nucleation and ice-crystal growth. The specimens were prepared by putting 2 μl of the sample between two aluminum planchettes with a gap of 100 μm between them. The specimen was then vitrified by liquid-nitrogen jet at 2100 bar for 360 msec in EM ICE system (Leica Microsystems, Wetzlar, Germany).

### 4.2. Production and characterization of pure curcumin and dasatinib particles

#### 4.2.1. Production of pure curcumin and dasatinib nanoparticles

Pure additive-free curcumin and DAS nanoparticles for *in vitro* and *in vivo* studies, respectively, were produced by using a microfluidic system with a Y-shape geometry that was designed and fabricated in our laboratory [67]. For this, curcumin (Alfa Aesar™, Haverhill, MA, USA) was initially dissolved in acetone (1 mL, final drug concentration was 0.2, Gadot, Netanya, Israel), whereas DAS (Carbosynth Ltd., Compton, UK) was initially dissolved in ethanol (Bio-Lab Ltd.). Then, the solution of each drug and water were injected by two syringe infusion pumps (SYP-01, MRC Ltd., Holon, Israel) into the channels of the microfluidic device and mixed rapidly in the intersection point at RT to produce the nanoparticles. The nanoparticles were collected in a 15 mL tube with a magnetic stirrer that stirred the nanosuspension at a constant stirring rate of 100 RPM. Acetone and ethanol were chosen as solvent since they dissolve very well the two compounds, are regarded as relatively safe [85] and are easy to eliminate by evaporation under ambient conditions. The size of the particles was controlled by adjusting the overall flow rate of each phase and the solvent and anti-solvent volume ratio that were finally set at 0.2:2.0 mL min^−1^ and 1:10 v/v, respectively, for both compounds. Once the precipitation process was completed, the nanosuspension was immediately frozen in liquid nitrogen and freeze-dried (Labconco Free Zone 4.5 plus L Benchtop Freeze Dry System) for further characterization. For penetration studies using particles of different size, curcumin particles in two sizes were used. To produce a curcumin suspension with a particle size of 1.2 μm, unprocessed curcumin was dispersed in water at a concentration of 0.2% w/v, vortexed (~1 min) and gently sonicated for 5 min to obtain a homogeneous suspension. For the preparation of curcumin particles with a size of 1.3 μm, several milligrams of raw curcumin were dispersed in water and filtered using a 2 μm filter paper (VWR^®^ Grade 410, qualitative, Strasbourg, France) under vacuum. The filtered particles were then left to dry at RT for 24 h, collected and redispersed in water (at a concentration of 0.2% w/v).

#### 4.2.2. Characterization of the nanoparticles

The D_h_ and PDI of the different nanoparticles were measured in a Zetasizer Nano-ZS (Malvern Instruments, Malvern, UK) at 25 °C with a 4 mW He–Ne laser (λ = 633 nm), a digital correlator ZEN3600 and Non-Invasive Back Scatter (NIBS^®^) technology at a scattering angle of 173° to the incident beam. DLS data were analyzed using CONTIN algorithms (Malvern Instruments). Values are expressed as Mean ± standard deviation (S.D.) and each measurement is a result of at least five runs. The S.D. of each size population, which is an expression of the peak width, was also determined. Differences among particle sizes were analyzed using one-way analysis of variance (ANOVA, significance level of 5%) with Bonferroni test (*p*<0.05).

The morphology of the pure curcumin and DAS nanoparticles was visualized by HR-SEM, carbon coating, acceleration voltage of 1–4 kV, Ultraplus). For HR-SEM, samples were dispersed in water and sprayed on top of a p-doped Si wafer <100> by introducing high-pressure N2, allowing the individual particles to be spread evenly on the wafer. Then, the wafer was attached to the grid using carbon-tape and additional tape was placed on its frame. Silver paint was applied to the corners of the frame prior to carbon coating.

Pure curcumin and DAS nanoparticles were analyzed by PXRD in an XRD diffractometer MiniFlex (Rigaku, Tokyo, Japan) under parallel-beam geometry at a speed rate of 6° min^−1^, θ–2θ range of 5–50 ° (with intervals of 0.01°) on a poly(methyl methacrylate) slide, at 25°C. Diffractograms of the pure drug nanoparticles were compared to those of the pristine drugs.

Thermal characterization of the raw curcumin and DAS and their corresponding nanoparticles were performed by DSC (2 STARe system equipped with a simultaneous thermal analyzer, STARe Software V13 and intra-cooler Huber TC100, Metter Toledo, Schwerzenbach, Switzerland). For this, samples (5–10 mg) sealed in 40 μL-Al crucible pans (Mettler Toledo) were heated from 25 to 325 °C at a heating rate of 10 °C min^−1^ under N2 gas flow (20 mL min^−1^) and Indium was used as a standard.

#### 4.3. Particle penetration studies *in vitro*

For particle penetration studies *in vitro*, we used H-CL mucin hydrogels (G’ = 200 Pa) and (i) curcumin particles of D_h_ of 200 nm, and 1.2 and 1.33 μm at a concnetration of 35 μg mL^−1^ and (ii) concentrations of 18, 35 and 71 μg mL^−1^ with a D_h_ of 200 nm. To produce pure curcumin nanoparticle concentrations of 18, 35 and 71 μg mL^−1^, we prepared a stock nanoparticle dispersion (concentration of 0.2 mg mL^−1^), evenly dispersed 50, 100 or 200 μL of this dispersion onto each mucin hydrogel (surface area of 1.9 cm^2^), and adjusted the water volume poured on top of each hydrogel to reach a total volume of 573 μL and produce a 3 mm-height water column; a water volume of 523, 473 and 373 μL was poured onto the hydrogels for 18, 35 and 71 μg mL^−1^ samples, respectively. To assess the effect of the crosslinking density of mucin hydrogels on particle penetration, we used H-CL, M-CL and L-CL hydrogels and pure curcumin nanoparticles with a D_h_ of 200 nm and at a concentration of 35 μg mL^−1^. Once the particles were dispersed onto the hydrogel and the 3-mm water colum produced, the system was incubated at RT. After 5, 10, 15, 20, 40, and 60 min, the supernatant above the hydrogel was carefully drawn-out and the hydrogel surface washed to ensure the recovery of 100% of the curcumin particles that did not penetrate the hydrogel. Water aliquots containing curcumin particles were frozen in liquid nitrogen, freeze-dried, re-dissolved in methanol and the concentration measured by fluorescence spectrophotometry (Fluoroskan^®^Ascent, ThermoFisher Scientific Oy) at *λ_ex_* of 438 nm and *λ_em_* of 538 nm and interpolated in a calibration curve of curcumin in methanol in the 0.25-10 μg mL^−1^ range to indirectly quantify the percentage of curcumin particles that penetrated the hydrogel by calculating the difference. Particle-free hydrogels subjected to the water column were used as control and processed following the same protocol. In each set of experiments, only one parameter, namely particle size, particle concentration, or crosslinking density of the hydrogel, was changed while keeping the others fixed to study the effect of each parameter separately.

#### 4.4. Comparative oral pharmacokinetics in rats

The oral PK of the unprocessed and nanonized DAS were assessed in albino rats (Sprague Dawley, Envigo, Jerusalem, Israel) following the approval of the preclinical protocol by the Committee of Animal Welfare of Technion (#IL-035-03-18, #IL-013-01-22). For this, animals (weight of 183-271 g, average of 230 ± 34 g) were divided into three groups and trained several days prior to the experiment to administer 9el gelatin capsules (Torpac, Inc., Fairfield, NJ, USA) and drink water (pH 2.8-3.2) from a 1 mL syringe to speed up the disintegration of the gelatin capsules. Then, formulations containing the exact drug dose based on the individual weight of each rat were weighed and loaded into the gelatin capsules before administration; The capsules were gently introduced into the mouth of fasted rats (12 h) using a metallic plunger (Torpac, Inc.). The capsule containing the formulation was accommodated in the cheek, minimizing animal discomfort (based on our previous experience [86]) and the animals received 1 mL of water using a plastic syringe. This procedure allows the fast disintegration of the capsule (less than 5 min) in the animal mouth and the smooth swallowing of the drug. Animals were placed in individual cages and followed up to ensure complete swallowing of the drug after the disintegration of the capsule. At predetermined time points post-administration, blood samples were extracted (250 μL), immediately centrifuged (3000 RPM, 4°C for 2 min, HERMLE Labortechnik GmbH, Wehingen, Germany) to separate plasma and frozen at −20°C until analysis. Then, the plasma was vortexed for 1 min and 30 μL of plasma were taken into a microtube. The proteins in plasma were precipitated by adding 100 μL HPLC grade acetonitrile (J.T. Baker, Avantor Performance Materials, Gliwice, Poland), vortexing and performing centrifugation (12,000 RPM, 4°C for 12 min). PK analysis was conducted by high-performance liquid chromatography (HPLC, Alliance, Waters Corp., Milford, MA, USA) with a separation module e2695 equipped with an XBridge C18 column (3.5 μm, 4.6×250 mm, Waters Corp., Dublin, Ireland) and UV-Vis detector (2998 Photoiodide Array UV/Vis 2D detector, W2998, Waters Corp.). The calculations of the following PK parameters were performed by non-compartmental analysis using PKSolver add-in program [87]: (i) C_max_, (ii) t_max_, (iii) AUC_0-∞_ and (iv) t_1/2_. The relative oral bioavailability (Fr) was calculated according to

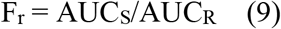

Where AUCS and AUCR are the AUC_0-∞_ of each sample and of unprocessed DAS (control), respectively. Results are expressed as mean ± coefficient of variation (CV %). Differences among groups were analyzed using one-way analysis of variance (ANOVA, significance level of 1%) with Bonferroni test.

## Acknowledgments

This research was funded by the Ministry of Science and Technology of Israel (MOST, Grant number 3-17339, “Alternatives to animal experiments”). A.S. thanks the support of the Tamara and Harry Handelsman Academic Chair. The partial support of the Russell Berrie Nanotechnology Institute (RBNI, Technion-Israel Institute of Technology) is also acknowledged.

