## Supplemental information for "An experimental and theoretical approach to understand the interaction between particles and mucosal tissues"

**Table S1. Hydrodynamic diameter ($D_h$) of pure curcumin nanoparticles produced by using a Y-shaped device over time, as measured by DLS at 25 °C.** Nanoparticles were produced using a Y-shaped device at a fixed drug solution concentration of 0.1% w/v and by setting flow rates of 0.2 and 2.0 mL $min^{-1}$ for the solvent and anti-solvent, respectively. A constant solvent/anti-solvent volume ratio of 1/10 was used.

| Time (Days) | $D_h$ (nm) – By intensity (± S.D.) [a] | Z-Average (nm) | S.D. (nm) [b] | PDI (± S.D.) |
|---|---|---|---|---|
| 0 | 200 (14) | 181 (10) | 66 | 0.09 (0.03) |
| 3 | 201 (4) | 156 (2) | 57 | 0.10 (0.01) |
| 6 | 200 (5) | 185 (6) | 61 | 0.10 (0.01) |

[a] $D_h$ are the intensity distribution values expressed as the average of five runs (n = 5) ± S.D., as determined by DLS.

[b] Standard deviation (S.D.) of each size population that is an expression of the peak width, as determined by DLS.

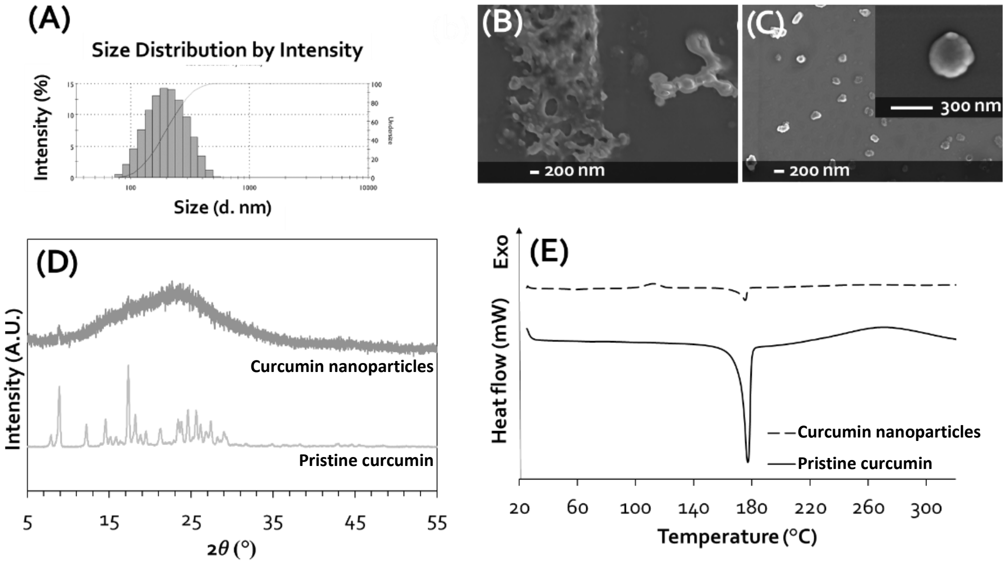


**Figure S1. Characterization of pure curcumin nanoparticles.** (A) Size distribution (by intensity), as measured by DLS. HR-SEM micrographs of (B) pristine curcumin and (C) pure curcumin nanoparticles. (D) X-ray diffraction patterns of pristine curcumin and pure curcumin nanoparticles. (E) DSC thermograms of pristine curcumin and pure curcumin nanoparticles.

**Table S2. Characterization of the hydrodynamic diameter ($D_h$) and the zeta-potential (Z-potential) of pure dasatinib and curcumin nanoparticles, as measured by DLS at 25 °C.** Nanoparticles were produced using a Y-shaped device at a fixed drug solution concentration of 0.1% w/v and by setting flow rates of 0.2 and 2.0 mL $min^{-1}$ for the solvent and anti-solvent, respectively. A constant solvent/anti-solvent volume ratio of 1/10 was used.

| Compound | $D_h$ – By intensity (nm) [a] (± S.D.) | S.D. (nm) [b] | Z-Average (nm) (± S.D.) | PDI (nm) (± S.D.) | Z-potential (mV) (± S.D.) |
|---|---|---|---|---|---|
| Curcumin | 200 (14) | 66 | 181 (10) | 0.09 (0.03) | -18 (0.4) |
| Dasatinib | 199 (6) | 63 | 123 (8) | 0.09 (0.02) | -18 (3) |

[a] $D_h$ are the intensity distribution values expressed as the average of five runs (n = 5) ± S.D., as determined by DLS.

[b] Standard deviation (S.D.) of each size population that is an expression of the peak width, as determined by DLS.

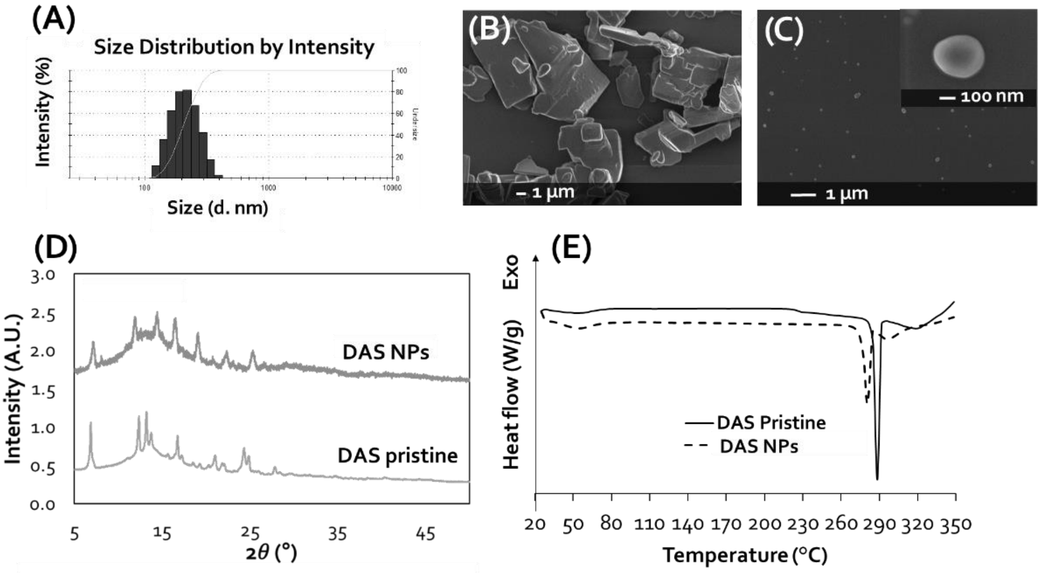


**Figure S2. Characterization of pure dasatinib nanoparticles.** (A) Size distribution (by intensity), as measured by DLS. (B,C) HR-SEM micrographs of (B) pristine dasatinib and (C) pure dasatinib nanoparticles. (D) X-ray diffraction patterns of pristine dasatinib and pure dasatinib nanoparticles. (E) DSC thermograms of pristine dasatinib and pure dasatinib nanoparticles.
